# Multilevel Plasticity and Altered Glycosylation Drive Aggressiveness in Hypoxic and Glucose-Deprived Bladder Cancer Cells

**DOI:** 10.1101/2023.10.21.561355

**Authors:** Andreia Peixoto, Dylan Ferreira, Andreia Miranda, Marta Relvas-Santos, Rui Freitas, Tim S. Veth, Andreia Brandão, Eduardo Ferreira, Paula Paulo, Marta Cardoso, Cristiana Gaiteiro, Sofia Cotton, Janine Soares, Luís Lima, Filipe Teixeira, Rita Ferreira, Carlos Palmeira, Albert J. R. Heck, Maria José Oliveira, André M. N. Silva, Lúcio Lara Santos, José Alexandre Ferreira

## Abstract

Bladder tumours with aggressive characteristics often present with microenvironmental niches marked by low oxygen levels (hypoxia) and limited glucose supply due to inadequate vascularization. The molecular mechanisms facilitating cellular adaptation to these stimuli remain largely elusive. Employing a multi-omics approach, we discovered that hypoxic and glucose-deprived cancer cells enter a quiescent state supported by mitophagy, fatty acid *β*-oxidation, and amino acid catabolism, concurrently enhancing their invasive capabilities. Reoxygenation and glucose restoration efficiently reversed cell quiescence without affecting cellular viability, highlighting significant molecular plasticity in adapting to microenvironmental challenges. Furthermore, cancer cells exhibited substantial perturbation of protein *O*-glycosylation, leading to simplified glycophenotypes with shorter glycosidic chains. Exploiting glycoengineered cell models, we established that immature glycosylation contributes to reduced cell proliferation and increased invasion. Our findings collectively indicate that hypoxia and glucose deprivation trigger cancer aggressiveness, reflecting an adaptive escape mechanism underpinned by altered metabolism and protein glycosylation, providing grounds for clinical intervention.

## Introduction

Bladder cancer (BLCA) remains one of the deadliest malignancies of the genitourinary tract due to high intra and inter-tumoral molecular heterogeneity^1^. This has delayed a more comprehensive understanding on tumour spatiotemporal status and affected the efficiency of precise clinical interventions. While genetic alterations are considered primary causes of cancer development, downstream phenotypic changes induced by the tumour microenvironment are amongst the driving forces of progression and dissemination. The generation of hypoxic niches characterized by decreased oxygen availability (≤2% O_2_) is a microenvironment hallmark of solid tumours^2^. Not surprisingly, the presence of hypoxic regions is a pivotal independent poor prognosis factor in several cancers, including urothelial carcinomas^3^.

Uncontrolled tumour cell proliferation supported by avid glucose consumption and glycolysis in the presence of oxygen and fully functioning mitochondria (the Warburg effect) is a common feature of solid tumours^4^. Rapid proliferation is frequently accompanied by flawed neoangiogenesis, resulting in suboptimal oxygen and nutrients supply to cancer cells in the periphery of blood vessels. Poor vascularization and competition for nutrients requires constant metabolic remodelling and exploitation of alternative survival strategies by cancer cells. While many tumour cells faced with suboptimal growth conditions undergo programmed cell death and necrosis, some subpopulations show tremendous molecular plasticity to adapt to hypoxic and nutrient deprived microenvironments^5^. Low oxygen levels induce metabolic rewiring towards anaerobic glycolysis and microenvironment acidification, while contributing to the maintenance of cancer stem cells and acquisition of epithelial-to-mesenchymal transition (EMT) traits, decisively dictating tumour fate^6,7^. Moreover, slow dividing cancer cells in hypoxic regions can escape many cytotoxic drugs targeting rapidly dividing cells, also being sufficiently shielded from many other therapeutic agents compared to the tumour bulk^8^. Although the mechanisms of cellular adaptation to hypoxia are already well known, few studies have interrogated how cancer cells can also withstand low levels of glucose. Furthermore, there is little knowledge on the underlying molecular alterations occurring at the cell surface, which dictate poor prognosis and may be easily targeted in theragnostic interventions.

The cell surface is densely covered by a layer of complex glycans (glycocalyx), which result from the concerted activity of a wide variety of glycosyltransferases, glycosidases, and sugar nucleotide transporters across the secretory pathways^9,10^. Moreover, glycosylation reflects microenvironmental stimuli in response to alterations in glycogenes expression and metabolic imbalances^11,12^. These alterations influence activation of oncogenic signalling, induction of immune tolerance, migration, cell-cell, and cell-matrix adhesion^11^. However, how hypoxia and glucose restriction shape the cancer glycome and its underlying functional implications remains mostly unaddressed.

In this study, we employed a multi-omics approach combining transcriptomics, metabolomics, glycomics, and phosphoproteomics to characterize in-depth the molecular plasticity of BLCA cells under hypoxia and low glucose and its contribution to aggressiveness. Having observed significant glycome remodelling in response to these stressors, we have conducted functional glycomics studies supported by a library of well-characterized glycoengineered cell models to determine how altered glycosylation impacts on BLCA progression. Important insights were generated to understand how BLCA adapts to oxygen and nutrient deprivation, aiming to identify more aggressive BLCA subpopulations envisaging future clinical interventions.

## Results

### Hypoxia and Low Glucose induce BLCA aggressiveness

To better understand the molecular adaptability of BLCA cells to hypoxia and low glucose, four BLCA cell models reflecting different grades of the disease (RT4-grade 1, 5637-grade 2, T24- grade 3, HT1197-grade 4) were cultured under low oxygen (0.1% O_2_) and reduced glucose (≤10%) levels. This was expected to mimic microenvironmental conditions encountered by cells growing far apart from blood vessels^13^. Cells responded rapidly to these challenges, stabilizing the hypoxia-inducible factor HIF-1α, which became more pronounced in the absence of glucose, supporting HIF-1α pivotal role in adaptive responses to microenvironmental stress (**Fig. 1a**). Notably, HIF- 1α was also increased in 5637 cells grown under atmospheric oxygen tension but with very low glucose (Normoxia-Glc), reinforcing the existence of a non-canonical regulation of HIF-1α stabilization regardless of oxygen availability^14^. The exception was the T24 cell line, which showed modest HIF-1α increase under stress; even though nuclear accumulation was notorious (**Fig.5a**). This was accompanied by an increase in lactate levels, that were rapidly extruded to the extracellular space under low oxygen (Hypoxia), suggesting the adoption of anaerobic glycolysis as main bioenergetic pathway and capacity to maintain intracellular homeostasis^7^ (**Fig. 1b**). Low glucose (Normoxia-Glc) also increased lactate as a result of aerobic glycolysis, except for HT1197 cells. However, it remained in the intracellular compartment, suggesting that low oxygen may be critical for activating extrusion mechanisms. Under these conditions, lactate may be converted into pyruvate to fuel the Krebs cycle. The combined effect of hypoxia and low glucose (Hypoxia-Glc) lowered lactate close to vestigial levels, strongly supporting the activation of an alternative energy-producing pathway to glycolysis.

**Fig. 1.**
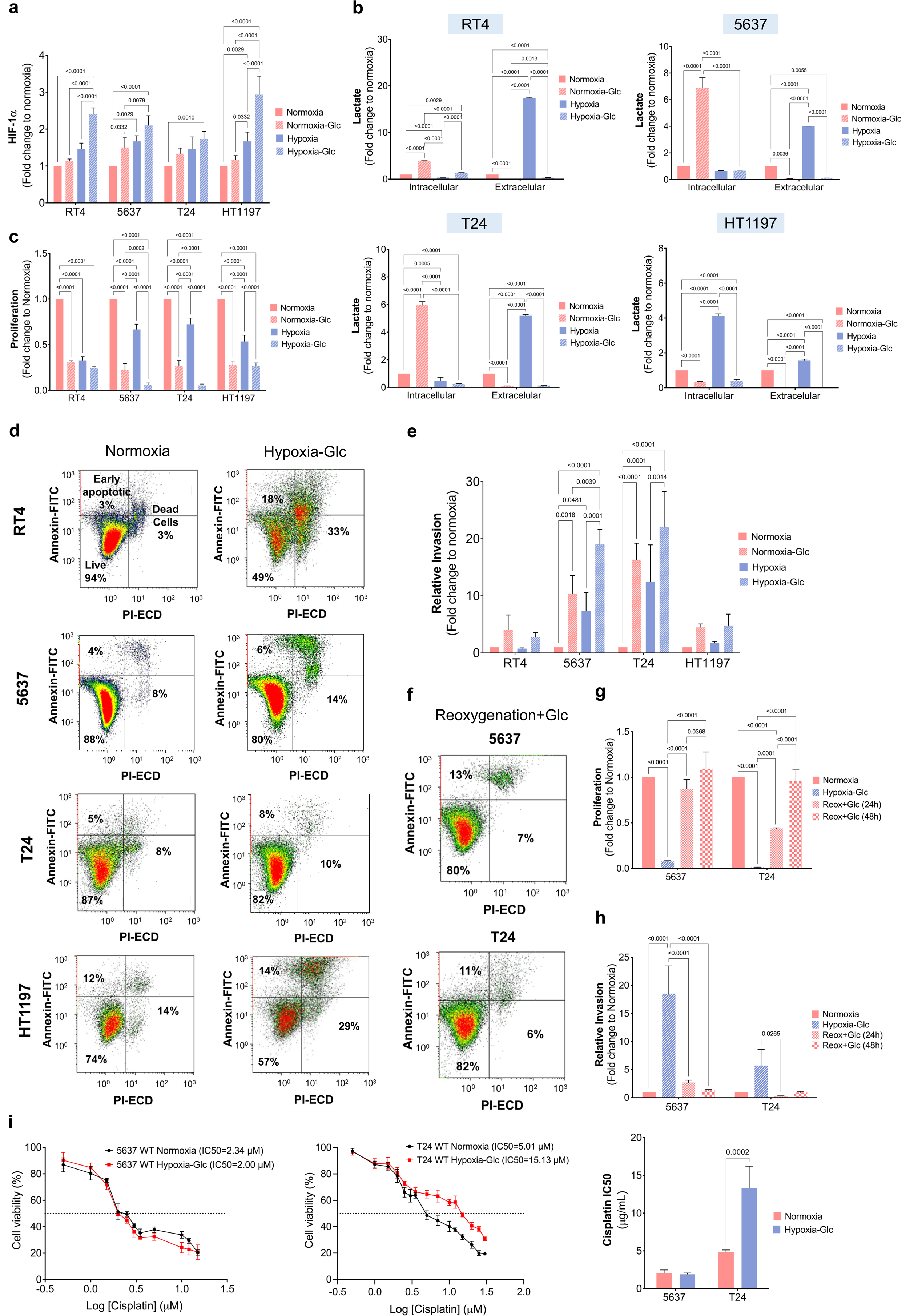
BLCA cells exhibit remarkable tolerance to hypoxia and low glucose, adopting a quasi-quiescent and more aggressive invasive behaviour. **a.** Hypoxia or low glucose (Normoxia -Glc) significantly upregulate HIF-1α expression in BLCA cell lines, which is further enhanced when combined (Hypoxia-Glc). **b.** BLCA cells cultured in hypoxia and low glucose produce residual levels of lactate. Individually, these stressors induce the production of lactate. **c.** Hypoxia and low glucose significantly suppress cell proliferation. Individually, low oxygen or low glucose inhibit cell proliferation. The combination of these stressors further exacerbates this effect in all cell lines. **d.** BLCA cells maintain their viability under hypoxia and low glucose. The combined environmental stress from hypoxia and low glucose does not significantly impact the viability of 5637 and T24 cells. RT4 and HT1197 cells exhibit a 50% reduction in viability under these conditions, suggesting a limited adaptive capacity. **e.** BLCA cells display increased invasiveness under hypoxia or low glucose. This is significantly potentiated when both stimuli are combined. f. BLCA cells demonstrate remarkable adaptability to microenvironmental changes with minimal impact on cell viability. Restoring oxygen and glucose levels does not affect cell viability, underscoring the high plasticity of these cells to endure drastic microenvironmental changes. **g.** BLCA cells restore basal proliferation after 48h of reoxygenation with glucose restoration. Both 5637 and T24 cells regain proliferative capacity, fully reinstating proliferation after 48h, highlighting their plasticity in responding to microenvironmental challenges. **h.** After 24h of reoxygenation with glucose restoration, BLCA cells exhibit a significant reduction in invasion, which is fully restored under normoxia after 48h. **i.** Hypoxia and low glucose increase T24 cells’ resistance to cisplatin across a wide range of concentrations, including its IC50, while 5637 cells remain unchanged. Error bars represent mean ± SD for three independent experiments. One-way ANOVA followed by Tukey’s multiple comparison test and the Mann-Whitney test were used for statistical analysis.

Concomitantly, we observed a striking decline in cell proliferation (approximately 1.5-fold) in all cell lines under hypoxia, which was significantly enhanced upon glucose suppression (**Fig. 1c**). Cells were arrested in late S phase (S/G2 transition), most likely due to depletion of the substrates required for DNA synthesis, as it was later confirmed by metabolomics (**Supplementary Fig. 1**). Notably, viability of 5637 and T24 cells was not affected after 24h of microenvironmental stress, as highlighted by little changes in the percentage of apoptotic and pre-apoptotic cells in comparison to normoxia (**Fig. 1d**). In contrast, RT4 and HT1197 cells viability was decreased by approximately 50%. Necrosis was not observed. Also, 5637 and T24 were able to withstand this stress for at least 72h without loss of cell viability. After 24h, 5637 and T24 cells responded to oxygen or glucose shortage by increasing invasion of matrigel *in vitro* (**Fig. 1e**). Strikingly, their combination (Hypoxia-Glc) enhanced further invasion, highlighting the pivotal role played by both stressors. Finally, reoxygenation and access to glucose restored proliferation after 24h and induced a massive drop in invasion without inducing apoptosis, suggesting little oxidative stress from drastic alterations in the microenvironment (**Figs. 1f-h**). Collectively, this demonstrated that certain BLCA cell subpopulations are well capable of accommodating hypoxia-induced stress, while concomitantly acquiring more aggressive and motile phenotypes. Overall, our results suggest that oxygen and glucose levels act as an on-off switch for proliferation and invasion. Interestingly, resistance to cancer cell death, proliferation decline, and activation of invasion traits have been closely linked to lactic acidosis as result of either hypoxia or glucose shortage^7^. Our observations support the adoption of similar behaviours in the absence of lactate. Finally, we addressed 5637 and T24 tolerance to cisplatin, generally used in the clinics against less proliferative tumour cells. Under stress, BLCA cells either maintained or significantly increased tolerance to cisplatin as observed for T24 cells (**Fig. 1i**), suggesting the adoption of cell chemoresistance mechanisms.

### Stress-induced transcriptome rewiring supports cancer aggressiveness and metabolic reprogramming

The 5637 and T24 BLCA cells showed markedly different transcriptomes, but common responses to hypoxia and glucose shortage (**Fig. 2a**). A total of 4,044 genes were differentially expressed in response to stress (1,722 upregulated, 2,322 downregulated; **Fig. 2b and Supplementary Table 1**), thus supporting significant transcriptome remodelling. Under hypoxia and glucose deprivation, cells activated stress-related genes driving more undifferentiated (*KRT17*)^15^, poorly proliferative (*PPP1R15A/GADD34*^16^), and cell death resistant (*PPP1R15A/GADD34*^17^, *DDIT4*^18^, *PFKFB3*^19^) phenotypes (**Fig. 2c**). Furthermore, upregulation of *DDIT4*, *HK2*, *PFKFB3*, and *PPP1R15A/GADD34* supported the activation of autophagic events^20–23^. In particular, *PPP1R15A/GADD34* plays a key role sustaining autophagy during starvation, thus enabling lysosomal biogenesis and a sustained autophagic flux^23^, whereas *DDIT4* has been linked to adaptive survival mechanisms permitting cells to resist metabolic stress^24^. Finally, we observed significant downregulation of genes linked to lipogenesis (*ELOVL6*)^25^, as well as both proliferation and positive regulators of differentiation (*FOSB*, *CSF3/G-CSF*)^26–28^.

**Fig. 2.**
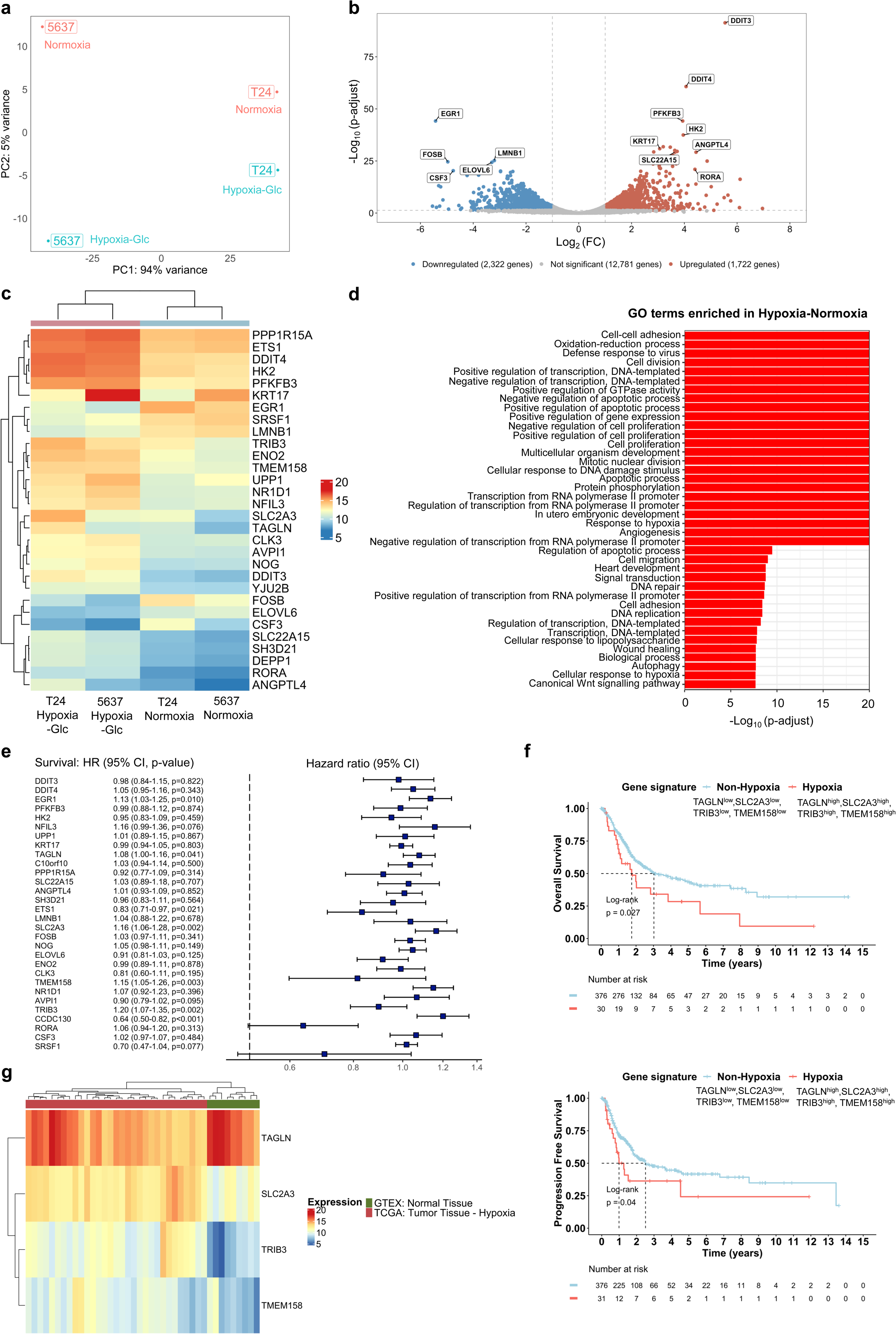
BLCA cell lines under hypoxia and low glucose experience profound transcriptome remodeling, linked to the acquisition of more aggressive phenotypes, which is supported by the poor prognosis observed in TCGA-BLCA patients. **a.** BLCA cell lines under hypoxia and low glucose display distinct transcriptomes but share common responses to these conditions. PCA for transcriptomics data reveals that PC1 (94% variance) primarily distinguishes differences between cell lines, while PC2 (5% variance) highlights marked changes between normoxic and stressed cells. **b.** Volcano plot showcases global transcriptional changes between normoxia and hypoxia plus low glucose. Exposure to these stressors alters the expression of 4,044 genes (1,722 upregulated, 2,322 downregulated), indicating significant transcriptome remodeling. **c.** Bi-clustering heatmap of the top 30 differentially expressed genes illustrates co-regulation under stress, supporting proliferation arrest, resistance to cell death, and invasion. Heatmap plots log2 transformed expression values of genes in samples. **d.** Enrichment analysis of GO terms for differentially expressed genes reveals alterations in key pathways associated with cell-cell adhesion, cell proliferation, and resistance to cell death. **e.** Prognostic evaluation identifies a hypoxia and glucose deprivation-linked four-gene signature (*TAGLN*^high^; *SLC2A3*^high^; *TRIB3*^high^; *TMEM158*^high^). Univariate Cox regression analysis of the top 30 differentially expressed genes identifies seven genes associated with OS. Higher expression levels of four genes, upregulated under hypoxia and low glucose, significantly correlate with poor OS, constituting a stress signature. **f.** Validation of the prognosis significance of the hypoxia-related four-gene signature in BLCA patients from TCGA. Kaplan-Meier curves of OS and PFS show significantly worse clinical outcomes for patients displaying the stress-related gene signature compared to the remaining patients in the cohort. **g.** Bi-clustering heatmap showing the association between the stress-related signature and bladder tumours. Heatmap plots log2 transformed expression values of the four hypoxia-related differentially expressed genes, showing clear differentiation between cancer and healthy bladder samples.

Furthermore, hypoxia and low glucose levels negatively regulated tumour suppressors (*EGR1*)^29^ and activated genes linked to invasive/migratory capacity (*ETS1*^30^, DDIT4^31^, TAGLN^32^, TMEM158^33^). Collectively, BLCA cells showed remarkable transcriptomic adaptability to microenvironmental stress supporting increased cancer aggressiveness. Genes involved in cell-cell adhesion, cell proliferation, programmed cell death, DNA damage, metabolism reprogramming, and oxidation-reduction processes (**Fig. 2d**) were triggered, in accordance with the observed functional alterations (**Fig. 1**). The transcriptome of bladder tumours (n>400 cases) showed significant correlation with hypoxia-related genes observed in stressed cell lines (**Supplementary Fig. 2**). Moreover, a distinct hypoxic phenotype (characterized by *TAGLN*, *SLC2A3*, *TRIB3*, and *TMEM158* upregulation) was identified in more aggressive tumours and associated with a worse prognosis (**Figs. 2e and f**). Importantly, this hypoxic phenotype was discriminative of tumours when compared to healthy tissues (**Fig. 2g**).

### Microenvironmental stress induces catabolic metabolism

To gain more insights on the metabolic reprogramming induced by hypoxia and glucose deprivation, we performed a mass spectrometry-based untargeted metabolomics study on 5637 and T24 cells (**Supplementary Table 2**). As highlighted by **Fig. 3a**, T24 and 5637 cells present distinct metabolic fingerprints under normoxia, in line with the different molecular backgrounds observed at the transcriptome level (**Fig. 2**). However, similar responses were observed facing low oxygen and glucose, characterized by a statistically significant reduction in the levels of 85 metabolites and increments in 8 species. Increased metabolites included several fatty acid-carnitine derivatives, while nucleotide sugar donors (UDP-Glc and UDP-GalNAc), citric acid, and gluconic acid were substantially decreased **(Figs. 3b and c).** High levels of fatty acid-carnitine derivatives (pentadecanoylcarninite, L-palmitoylcarnitine, heptadecanoylcarnitine, stearoylcarnitine, arachidylcarnitine; **Figs. 3b and c**) support active translocation of long-chain fatty acids across the inner mitochondrial membrane for subsequent *β*-oxidation. Interestingly, increased levels of these metabolites have been observed in the urine of advanced stage BLCA patients^34,35^, reinforcing the close link between this metabolic phenotype and cancer aggressiveness. In addition, stressed cells showed decreased amounts of several lysophosphatidylcholines and lysophosphatidylethanolamines, which is consistent with the mobilization of lipids for mitochondrial *β*-oxidation^36^ **(Fig. 3c)**, and in perfect agreement with the lipolytic phenotype highlighted by transcriptomics Interestingly, carnitine levels were also conserved to maintain downstream lipid *β*-oxidation at the expenses of L-lysine and L-methionine degradation (**Fig. 3c- e**). On the other hand, stressed cells showed significant amounts of Krebs cycle intermediates, namely citrate, but also oxoglutarate, oxaloacetate, malate, and fumarate. Interestingly, lower citrate in cancer cells has been described to favour resistance to apoptosis and cellular dedifferentiation^37^, thus in agreement with functional and transcriptomics studies. Of note, we also observed amino acids consumption (lysine, valine, leucine, isoleucine; **Fig. 3d**) to support energy requirements. A joint pathway analysis, combining transcriptomics and metabolomics confirmed the molecular rewiring of challenged cells towards a catabolic state (**Fig. 3e**). It further highlighted changes in glycolysis and gluconeogenesis, in agreement with the adoption of a lipolytic rather than a glycolytic metabolism.

**Fig. 3.**
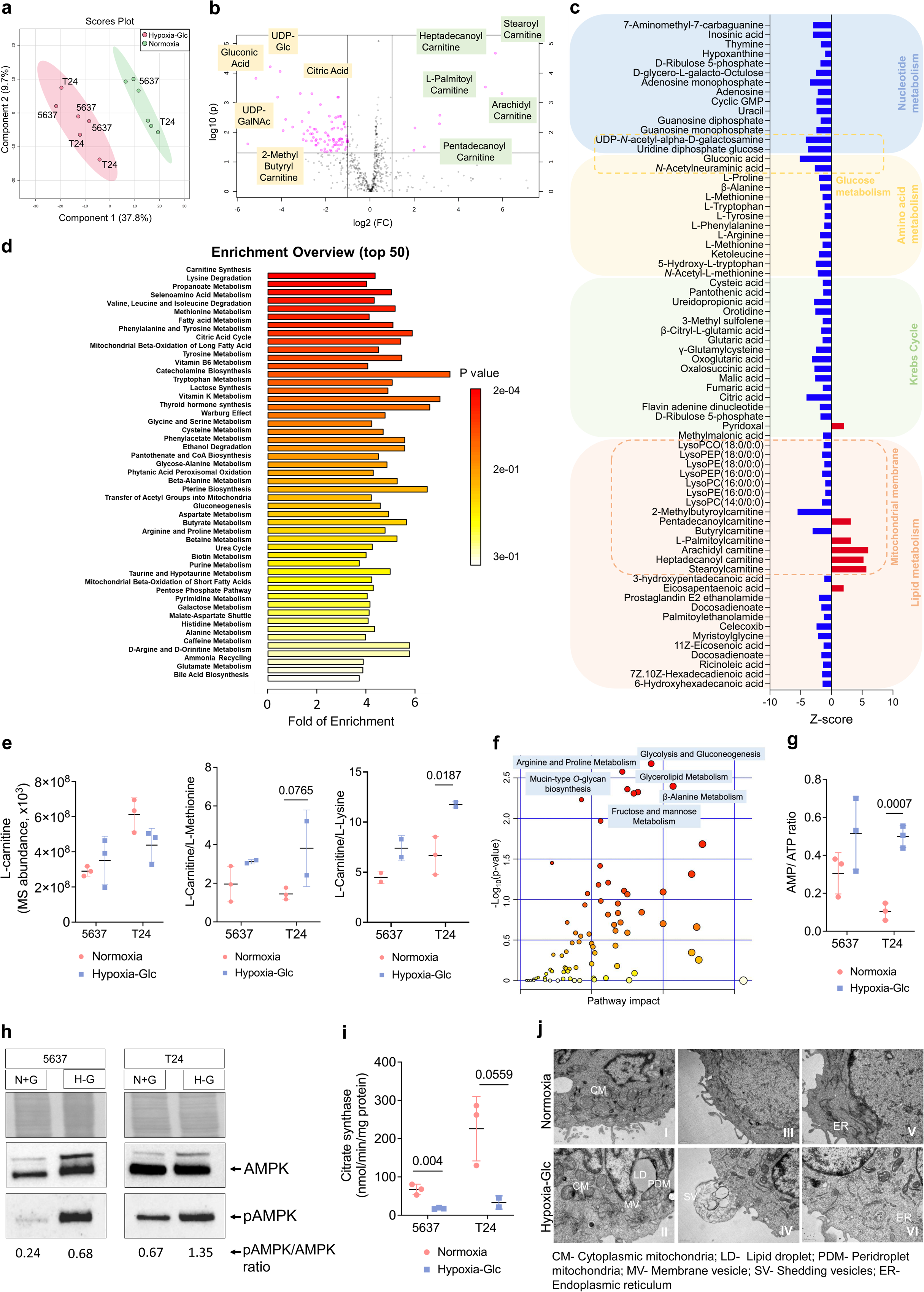
Hypoxia and low glucose shift BLCA cell metabolism from glycolytic to lipolytic, increasing lipid droplet formation and reducing the number of active mitochondria. a, b, c. PLS-DA analysis reveals similar metabolic responses in 5637 and T24 cells under microenvironmental stress **(a).** Volcano plot highlights significant metabolome alterations in response to hypoxia and low glucose **(b).** Downregulated metabolites include UDP-Glc, UDP-GalNAc, gluconic acid, and citric acid, while increased metabolites indicate active fatty acid transport and *β*-oxidation **(c).** Significant reduction in key metabolites linked to nucleotide, amino acids, Krebs cycle, and lipid metabolism was observed, consistent with catabolic metabolism. Exception is the accumulation of long fatty acid acylcarnitine for transfer across the inner mitochondrial membrane for *β*-oxidation. **d.** Pathway enrichment analysis supports fatty acid *β*- oxidation as the primary bioenergetic pathway in stressed cells. Key metabolic pathways, including carnitine biosynthesis and lysine/methionine degradation, contribute to fatty acid β-oxidation. **e.** Hypoxia and low glucose induce lysine and methionine degradation to support acylcarnitine biosynthesis and lipid *β*-oxidation. **f.** Joint pathway analysis incorporating transcriptomics and metabolomics studies supports changes from glycolytic to lipolytic metabolism, impacting nucleotides and sugars biosynthesis, including *O*-GalNAc glycans and protein *O*-glycosylation. **g, h**. Hypoxia and low glucose increase AMP/ATP ratio (**g**) and activate AMPK by phosphorylation (**h**), indicating impaired oxidative phosphorylation and potential catabolic processes, including mitophagy. **i.** Citrate synthase activity decreases under hypoxia and low glucose, suggesting a reduction in functional mitochondria. **k.** TEM analysis reveals major morphological changes, including compromised mitochondria, lipid droplets, peridroplet mitochondria, membrane vesicles, and increased shedding of vesicles, indicating membrane activity changes under stress. Error bars represent mean ± SD for three independent experiments. Mann-Whitney Test was used for statistical analysis.

Not surprisingly, we found higher AMP/ATP ratio in stressed cells (**Fig. 3g**), mainly driven by a significant decrease in ATP. This was accompanied by higher 5-AMP-activated protein kinase (AMPK) phosphorylation (**Fig. 3h**), frequently observed in more aggressive bladder tumors^38^. These findings were both consistent with the adoption of catabolic processes and supported the occurrence of mitophagy. Accordingly, microenvironment challenged cells displayed decreased citrate synthase activity (**Fig. 3i**) and higher number of autophagosomes (**Supplementary Fig. 3a**). Autophagy in hypoxic cells was confirmed by dedicated immunoassays (**Supplementary Fig. 3b**). Mitophagy was also later confirmed by TEM **(Fig. 3j)**. In fact, microscopy evidenced a drastic decrease in the number of intact mitochondria, allied to evident mitophagy events (**Figs. 3j-I and II**), translated by outer mitochondrial membrane-associated degradation and matrix sectioning. Lipid droplet (LD)-associated mitochondria, also known as peridroplet mitochondria (PDM), were also observed under hypoxic conditions **(Fig. 3j-II)**. Vesicles shedding was evident **(Figs. 3j-III and IV)**, highlighting cellular communication events that should be carefully investigated in future studies. Finally, stressed cells showed considerably short and disorganized endoplasmic reticulum (ER) cisternae, contrasting with typically longer ER sections presented by cancer cells in normoxia **(Figs. 3j-V and VI)**, demonstrating prominent disorganization of secretory pathways where glycosylation occurs. Overall, cellular ultrastructure was significantly altered in ways that are consistent with the underlying molecular alterations.

### Cellular Signalling rewiring under stress supports aggressiveness

Cellular signalling rewiring was assessed by phosphoproteomics (**Supplementary Table 3**). Under stress, 5637 and T24 BLCA cells responded similarly **(Fig. 4a)**, exhibiting major alterations in relevant serine/threonine kinases and substrates linked to high motility and reduced cell adhesion, reduced cell proliferation, cell senescence, and mitochondrial reorganization (**Fig. 4b and c; Supplementary Table 3**). Moreover, cells exhibit clear survival adaptations translated by induction of autophagy, apoptosis evasion, cell-cycle arrest, krebs cycle inhibition, and cell-survival (**Fig. 4**), in full agreement with functional, transcriptome and metabolome reprogramming. Increased endocytosis was also a relevant feature of these cells, supporting major plasma membrane remodelling also observed by TEM. Notably, most significant alterations in phosphorylation (>2-fold change to normoxia) occurred in proteins involved in the AMPK, insulin, HIF-1α, EGFR tyrosine kinase and autophagy signalling pathways (**Fig. 4e**). The main altered phosphorylation sites under hypoxic conditions are AKT1 S473, AKT3 S472, PFKFB3 S461, CAMKK2 S511, PDHA1 S232, BAD S118, and PRKAB1 S108 (**Fig.4d; Supplementary Table 3**). AKT1 S473 and AKT3 S472 act synergically to protect cells from microenvironmental and cisplatin-induced apoptosis by phosphorylating downstream molecules as mTOR and glycogen synthase kinase-3. Notably, under stress, the preferable PRKCZ substrate is AKT3 whose activity was found significantly increased (**Fig. 4e**). PRKCZ has been implicated in maintaining high motility and reduced cell adhesion of metastatic cancer cells^39^, in cell cycle regulation^40^, and in mitochondrial reprogramming in senescent cells towards cell survival. Also, BAD apoptotic activity is attenuated by phosphorylation at S118, precluding binding to other Bcl-2 family members to evade apoptosis^41^. Furthermore, phosphorylated PDHA1 at S232 inactivates pyruvate dehydrogenase complex^42^, redirecting cells toward alternative energy pathways to Krebs cycle. Together with elevated phospho-PRKAB1 S108, this is consistent with the adoption of catabolic processes such as fatty acid *β*-oxidation and potentially mitophagy^43^ sustained by AMPK activation (**Fig. 4d and e**). A significant decrease in several forms of RPS6 (S240, S235, S244), RAF1 S621, ELAVL1 S202, and EIF4EBP1 T70 is also compatible with impaired cancer cell proliferation observed under stress^44–47^. These alterations are consistent with the adoption of more aggressive phenotypes by microenvironment challenged cells.

**Fig. 4.**
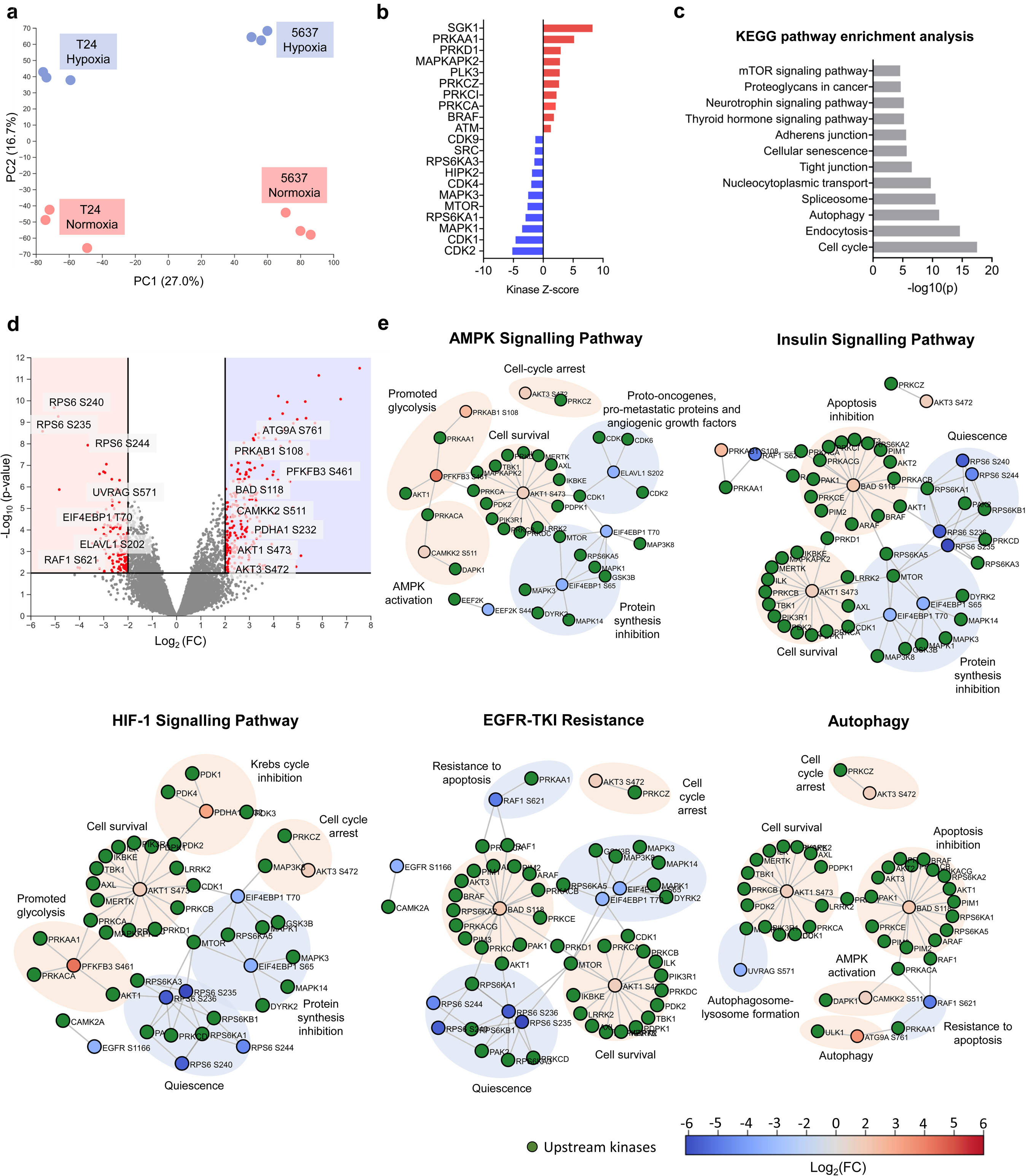
Hypoxia and low glucose induce major cell signalling rewiring, promoting aggressiveness via cell cycle arrest, quiescence, apoptosis resistance, glycolytic metabolism blockage, and autophagy. **a.** Despite cell-related differences, common features in oncogenic signalling are elicited by hypoxia and glucose deprivation, as shown by PCA analysis of phosphoproteome signatures. PC1 (27% variance) distinguishes cell-dependent differences, while PC2 (17% variance) relates to marked signalling changes linked to hypoxia and glucose deprivation. **b.** Kinase-Substrate Enrichment Analysis scores each kinase based on substrate phosphorylation, highlighting major alterations in stressed cells. Red indicates significantly activated kinases, while blue indicates significantly inhibited ones (z-score ≥ 2 or ≤2). **c.** KEGG pathway enrichment analysis of phosphoproteomics data shows significant alterations in cell signalling pathways supporting cell motility, cellular senescence, and autophagy under stress. **d.** Volcano plot showcases global cell signalling rewiring between normoxia and hypoxia plus low glucose, indicating the most significantly hyper or hypo-phosphorylated proteins and precise annotation of main phosphorylation sites. **e.** AMPK, insulin signalling, HIF-1α signalling, EGFR-TKI resistance, and autophagy are the most significantly altered pathways in stressed cells. Top-ranked signalling pathways with 2-fold change in stressed cells are presented, highlighting the most significant hyper or hypo-phosphorylated proteins and precise phosphorylation sites, along with main associated cellular functions.

### Stress induces an immature glycocalyx

The generalized disorganization of secretory organelles on stressed cells strongly suggests implications for the cells’ glycocalyx. In addition, **Fig. 3** supports major alterations in glycosylation pathways driven by changes in metabolites linked to glycosylation. This includes UDP-GalNAc, a key sugar nucleotide for the initiation of protein *O*-GalNAc glycosylation and Neu5Ac, a sialic acid capping different glycoconjugates (**Fig. 3c**). Data also suggests low flux through the hexosamine pathway, which impacts on UDP-GlcNAc biosynthesis required for glycosylation, even though this metabolite could not be detected by our approach. We further observed decreased UDP-Glc, which is a key precursor of UDP-Gal and UDP-GlcA needed for glycoproteins, proteoglycans, and glycolipids glycosylation. Finally, lower levels of gluconic acid and D-ribulose 5-phosphate as well as AMP, UMP, GMP, and GDP demonstrate alterations in the pentose phosphate pathway (**Figs. 3c and d**) required to generate nucleotides for nucleic acid synthesis but also glycosylation. Major inhibition of mucin-type *O*-GalNAc glycans biosynthesis as well as fructose and mannose metabolism required for glycosylation was also evident on the joint pathway analysis (**Fig. 3e**). In addition, downregulation of several glycogenes encoding polypeptide *N*-acetylgalactosaminyltransferases responsible for protein *O*-glycosylation initiation (**Supplementary Table 4**) was evident from transcriptome analysis. Downregulation of *C1GALT1C1*, encoding an essential chaperone for core 1 *O*-glycan T-synthase activity and further *O*-glycans elongation, was also observed. Interestingly, stressed cells also presented downregulation of key glycosyltransferases for early *N*-glycans processing in the ER, but also *N*- and *O*-glycans elongation in the Golgi. Finally, we observed downregulation of several glycogenes linked to proteoglycans biosynthesis (**Supplementary Table 4**). In summary, disorganization of secretory organelles and decreased sugar nucleotides, accompanied by a net downregulation of key glycogenes linked to glycosylation initiation and elongation, drive an immature glycophenotype.

Building on our previous studies linking immature glycosylation with BLCA aggressiveness, we devoted to the precise characterization of the *O*-GalNAc glycome of stressed cells. This was performed by exploiting the Tn mimetic benzyl-α-GalNAc as a scaffold for further *O*-chain elongation, providing a good assessment of *O*-glycosylation pathways fitness **(Fig. 5)**. According to **Fig. 5a**, low oxygen and glucose significantly reduced *O*-glycans synthesis in both cell lines (**Supplementary Fig. 4 and Table 5**). More detailed glycomic characterization in **Fig. 5c** showed that both cell lines abundantly express fucosylated (*m/z* 746.40; H1N1F1; type 3 H-antigen) and sialylated T (*m/z* 933.48; H1N1S1) antigens, also exhibiting several extended core 2 *O*-glycans of variable lengths, degrees of fucosylation and sialylation. Low amounts of shorter *O*-glycans such as core 3 (*m/z* 613.33; N2) and STn (*m/z* 729.38; N1S1) antigens could also be observed. The drastic reduction of oxygen and glucose significantly impacted the glycome of cells, inducing a simple cell glycophenotype characterized by an accumulation of few short-chain *O*-glycans without chain extension beyond core 1, including core 3 and, to less extent, mono- (*m/z* 933.48; H1N1S1) and di-sialylated (*m/z* 1294.65; H1N1S2) T antigens (**Figs. 5a-c**). Trace amounts of STn antigen could also be detected. Interestingly, no extension of core 3 was observed, reinforcing the inexorable expression of shorter structures by stressed cells. Furthermore, we found that the main driver for the lack of glycan extension was the reduction in glucose (**Fig. 5c**). Notably, BLCA cells regained the capacity to extend glycans after reoxygenation and reintroduction of glucose (**Fig. 5c**), demonstrating significant plasticity and that microenvironmental stressors act as an on-off switch for *O*-glycosylation. Complementary flow cytometry studies showed that the percentage of cells expressing T antigen decreased under hypoxia and glucose deprivation in comparison to normoxia, whereas the abundance of sialylated T antigens was significantly increased (**Fig. 5b**). Also, the percentage of Tn expressing cells increased under stress. Overall, these findings demonstrate that core 2 extension is repressed under hypoxia and glucose deprivation, leading to the accumulation of sialylated T and, to less extent, Tn antigens. Interestingly, we found similar patterns in a wider panel of cell lines, including RT4 and HT1197, but also oesophageal, gastric, and colorectal cancer cells and human monocyte-derived macrophages (**Supplementary Fig. 5 and Table 6**), suggesting common response patterns to stress. Notably, in these cells, core 2 was significantly decreased but not completely lost. Finally, we exposed BLCA cells to deferoxamine (DFX), an ion chelator known to stabilize HIF-1α expression and confirmed nuclear accumulation of HIF-1α (**Fig. 5a**). Upon exposure to DFX, both cell lines exhibited a significant decrease in the expression of *C1GALT1C1*, a crucial glycosyltransferase responsible for *O*-chain elongation beyond the Tn antigen. Additionally, in 5637 cells, *GCNT1*, which regulates core 2 biosynthesis, is downregulated, while T24 cells upregulate *ST3GAL1* **(Supplementary Fig. 6)**. These alterations could potentially contribute to a premature stop in glycans elongation. Interestingly, DFX did not induce significant changes in the glycosylation of both cell lines, indicating that the HIF-1α transcription factor might not exert a major influence on glycome remodelling, despite its role in regulating crucial *O*-glycogenes expression. Our observations also highlight the need for careful interpretation of the glycome based on glycogenes expression. Finally, seeking deeper insights on the relation between glycogenes expression and the glycophenotype, we re-analysed by RT-PCR the expression of a wide array of glycosyltransferases involved in the initial steps of *O*-glycans biosynthesis (**Fig. 5d**). Glucose suppression impacted more significantly on glycogenes expression than hypoxia, inducing significant overexpression of glycosyltransferases linked to *O*-glycans elongation, namely *C1GALT1*, *C1GALT1C1* (for T24 cells), *GCNT1,* and *B3GNT6*, but also extension and termination by fucosylation and/or sialylation (**Fig. 5d**). *ST3GAL1* overexpression was observed and may contribute to a premature stop in core 1 extension by *O*-3 sialylation, decreasing core 1 fucosylation. Interestingly, the combination of both hypoxia with low glucose impacted less on glycogenes remodelling than glucose suppression alone (normoxia-Glc). Notably, *B3GNT6* and *ST3GAL1* remained overexpressed (**Figs. 5d**). Downregulation of *C1GALT1* and *C1GALT1C1* were also observed (**Fig. 5d**), which may contribute to reduce core 1 synthesis and further elongation to core 2, impacting negatively on glycans extension. However, western blot analysis showed that, despite changes in glycogenes expressions, ST3Gal-I was the only main glycosyltransferase gatekeeper of *O*-glycans extension found elevated in stressed cells. (**Fig. 5e**). Also, while the focus was set on *O*-GalNAc glycosylation, we also characterized the *N*- glycome. Despite changes in *N*-glycogenes expression (**Supplementary Table 4)**, no major alterations could be observed in relation to the main types of *N*-glycans (oligommanose, paucimannose, complex; **Supplementary Table 7**).

**Fig. 5.**
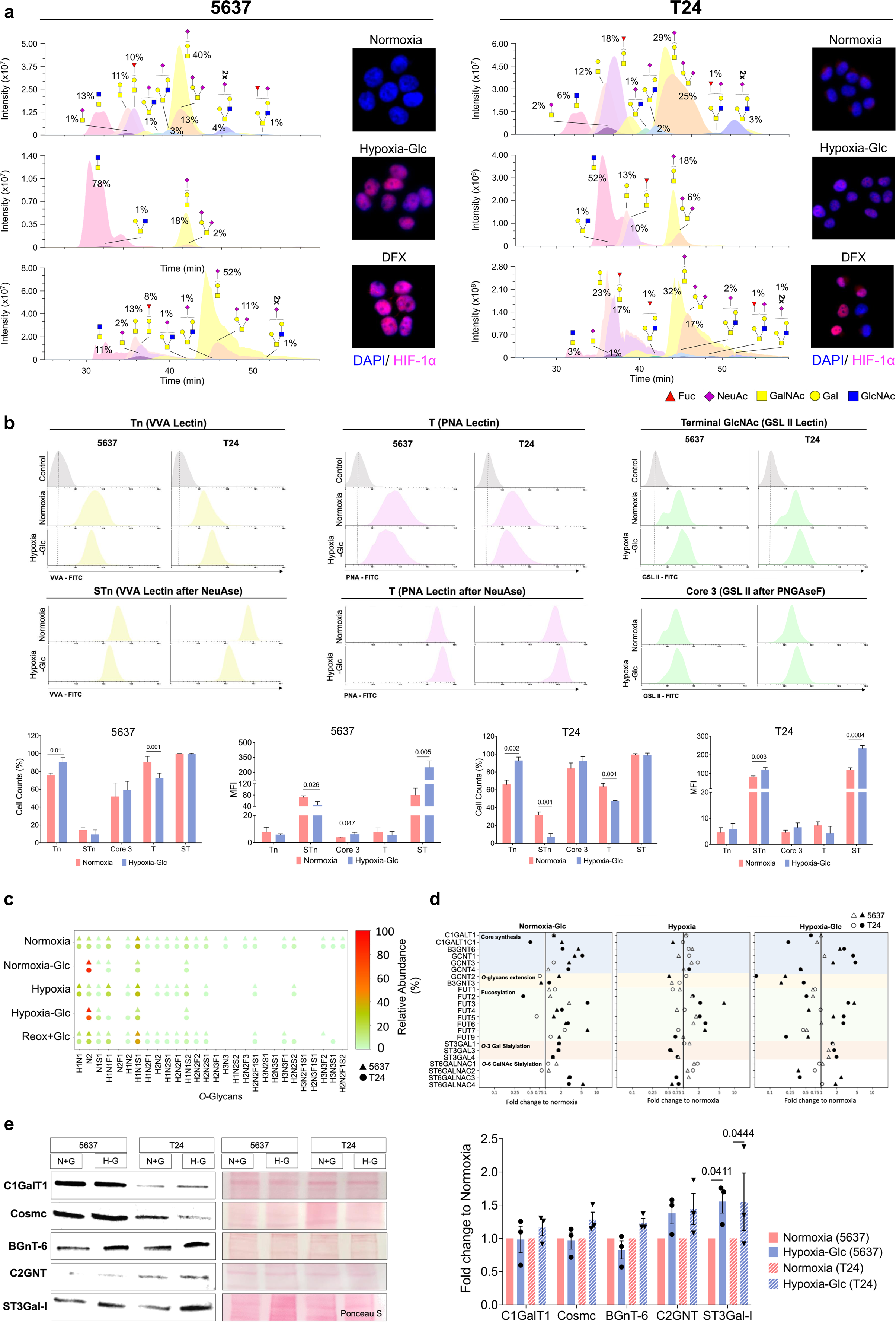
Hypoxia and low glucose impair *O*-glycans extension in BLCA, originating a simple cancer cell glycophenotype. **a.** BLCA cells exposed to hypoxia and low glucose exhibit reduced abundance, simpler, and shorter glycomes, lacking extensions beyond core 1 structures. nanoLC-MS/MS analysis shows that this glycophenotype is characterized by increased sialylated T antigens and abundant core 3, likely due to decreased typical core 1 and 2 structures. DFX-treated cells, stabilizing HIF-1α, show no significant alterations in the glycome, suggesting that changes observed in stressed cells are not driven by HIF-1α. **b.** Lectin affinity studies show significant upregulation of Tn and sialylated T antigens (recognized by PNA lectin after NeuAse digestion) under stress, in accordance with MS-based glycomics. Notably, core 3 *O*-glycans (evaluated by GSL II lectin after PNGase F digestion) remain unchanged, highlighting that cellular stress primarily suppresses core 1/2 *O*-glycans, rather than increasing core 3 *O*-glycans. **c.** Glucose suppression is the primary driver of glycome remodelling, which can be reversed by reoxygenation and restoration of glucose. **d.** Glycogene remodelling is primarily driven by the combined effects of hypoxia and glucose deprivation and leads to a premature halt in glycans extension beyond core 1. *C1GALT1C1*, necessary for core 1 biosynthesis, is downregulated, while *ST3GAL1, 3,* and *4* are overexpressed, increasing sialylated T antigens and inhibiting core 2 formation. Downregulation of *GCNT4* also contributes to core 2 inhibition. Interestingly, elevated *GCNT1* and *GCNT3* potentially counterbalances core 2 suppression **e.** Quantification of key enzymes involved in *O*- glycan elongation (C1GalT1; Cosmc; BGnT-6; C2GNT; ST3Gal-I) shows significant upregulation of ST3Gal-1 in stressed cells, consistent with transcriptomics. The others remain unchanged, indicating distinct regulation between glycogenes and glycosyltransferases under these conditions. Bold circles and triangles represent statistically significant changes in T24 and 5637 cell lines, respectively. Error bars represent mean ± SD for three independent experiments. Mann-Whitney Test was used for statistical analysis.

### Hypoxic bladder tumours share common molecular features with stressed cell lines

We then assessed the translational character of our observations in patient samples. A cohort of BLCA surgical tissue sections without prior history of adjuvant treatments (n=70), comprehending non-muscle (NMIBC; n=30) and muscle invasive (MIBC; n=40) tumours spanning all stages of the disease (Ta, T1, T2, T3, T4) was elected for this study. The tumours were screened for nuclear expression of HIF-1α and expression of the proliferation marker Ki-67 (HIF-1α^+^ Ki-67^low^ ; hypoxic phenotype). HIF-1α was detected in 10% of MIBC tumours, showing heterogeneous expression ranging from 20 to 60% of the tumour area (**Fig. 6a**). HIF-1α was not found in superficial lesions (**Fig. 6a**). Notably, 59% of studied MIBC were poorly proliferative, including the 4 hypoxic tumours, whereas only 15% of NMIBC presented a Ki-67 ^low^ phenotype. Also, HIF-1α-positive tumour areas did not express Ki-67, in agreement with the acquisition of quiescent phenotypes by hypoxic cells (**Fig. 6b**). However, no associations between the hypoxic phenotype and prognosis were found, as these were already highly aggressive lesions.

**Fig. 6.**
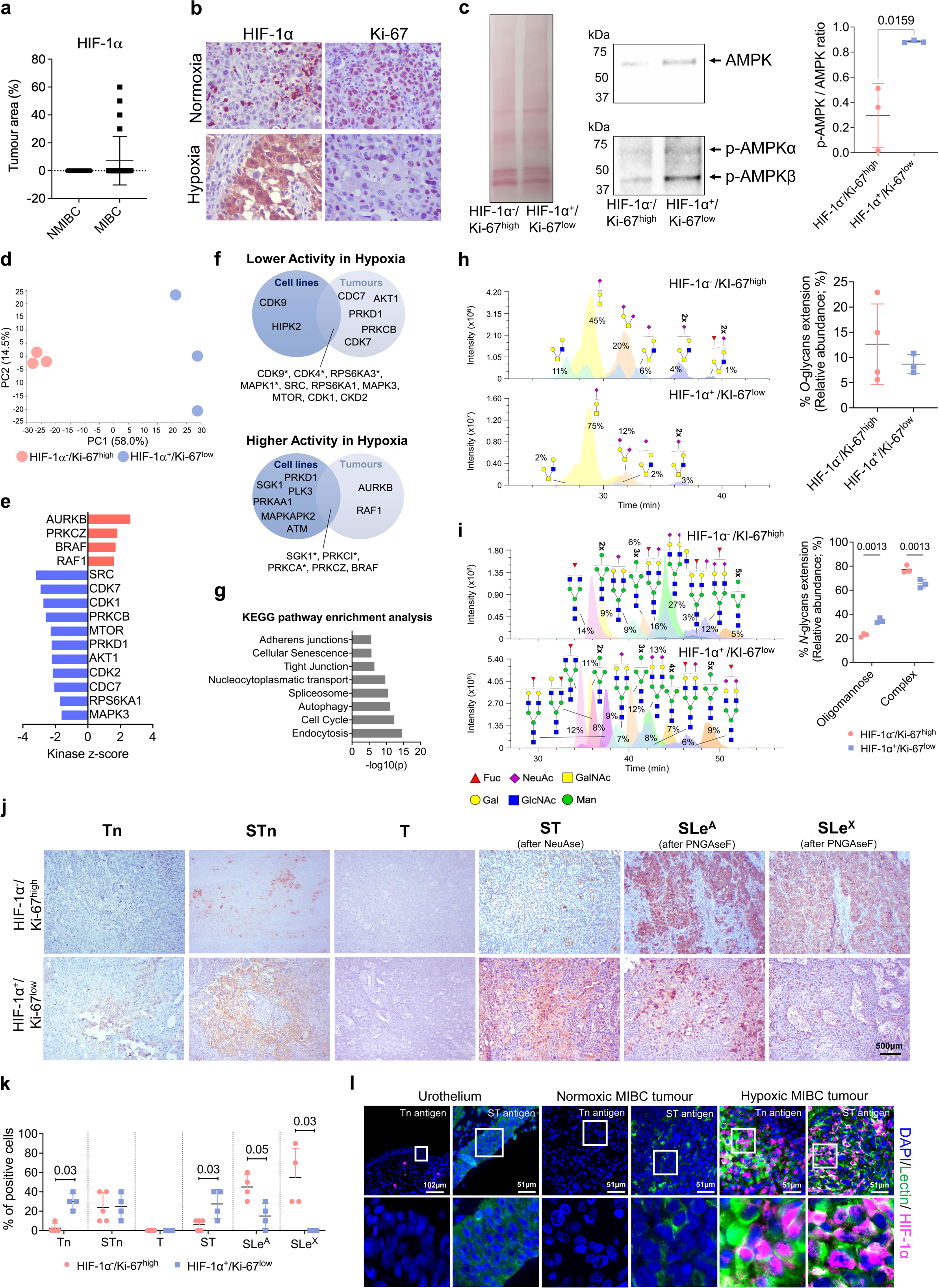
Hypoxic BLCA, characterized by high nuclear HIF-1α expression and low proliferation, shares malignant molecular features with hypoxic and glucose deprived cells *in vitro*, including simple glycophenotypes. a, b. Roughly 10% of MIBC tumours display a hypoxic fingerprint (HIF-1α^positive^/Ki-67^low^) that was not observed in NMIBC and most MIBC tumours (HIF-1α^negative^/Ki-67^high^), indicating a potential link to aggressiveness. **c.** Hypoxic tumours display significantly higher AMPK phosphorylation compared to proliferative cases, denoting a catabolic state. **d.** Hypoxic tumours show distinct cellular signalling pathway activation compared to proliferative tumours. PCA for phosphoproteomics data indicates that PC1 (58% variance) primarily separates hypoxic from proliferative tumours, while PC2 (15% variance) highlights marked differences among hypoxic tumours. **e.** Kinase-Substrate Enrichment Analysis supports major cell rewiring in hypoxic tumours. Kinases color-coded in red are significantly activated, while blue are significantly inactivated. **f.** Hypoxic tumours share common kinase activation patterns with stressed BLCA cells *in vitro*. **g.** KEGG pathway enrichment analysis indicates significant alterations in cell signalling pathways, promoting cell motility, cellular senescence, and autophagy in hypoxic tumours as found in stressed cells *in vitro*. **h.** Hypoxic tumours present simple *O*-glycophenotypes compared to proliferative tumours. nanoLC-MS/MS reveals more homogeneous *O*- glycome in hypoxic tumours with scarce core 2 glycans. **i.** Hypoxic tumours *N*-glycome is enriched for oligomannoses, whereas proliferative tumours are enriched for complex *N*-glycans. **j, k.** Hypoxic tumours show higher levels of Tn and sialylated T antigens and lower levels of sialylated Lewis antigens in *O*-glycans compared to proliferative tumours, reinforcing the primary suppression of *O*-glycan extension. **l.** In hypoxic tumours, Tn and sialylated T antigens co-localize with high HIF-1α. Normoxic, proliferative tumours lack HIF-1α and show low levels of sialylated T antigens and no Tn antigens. Healthy urothelium from non-cancerous individuals served as a negative control for HIF-1α, low Tn, and sialylated T antigens expression. Unpaired T-test and Mann-Whitney Test were used for statistical analysis.

We then elected for downstream molecular studies tumours showing diffuse and intense nuclear HIF-1α expressions (>50% of the tumour area). These tumours showed increased AMPK phosphorylation and pAMPK/AMPK ratios in comparison to highly proliferative lesions (**Fig. 6c**), consistent with a catabolic state. Subsequent phosphoproteomics highlighted marked differences between hypoxic and non-hypoxic tumours, and striking similarities between hypoxic tumours and cell lines in terms of kinase activation (**Figs. 6d and f; Supplementary Table 3**). Namely, we found increased PRKCZ and BRAF phosphorylation, contrasting with significantly lower SRC, CDK1, MTOR, CDK2, RPS6KA1, and MAPK3 activation (**Figs. 6e** and **f**). Likewise, altered pathways were also common, with emphasis on endocytosis, cell cycle arrest, senescence, autophagy, and invasion/cell motility (**Fig. 6g**). Furthermore, hypoxic tumours exhibited elevated expression of hypoxia-associated poor-prognosis markers *TMEM* and *SLC2A3* (**Fig. 2f and Supplementary Fig. 7**). Moreover, 75% of hypoxic tumours co-overexpress at least two out of four hypoxia-linked genes with prognostic significance (**Fig. 2f**), in contrast to 33% under normoxia, underscoring their molecular resemblance to stressed cells *in vitro*.

The glycome of hypoxic tumours was then characterized by mass spectrometry, revealing low levels of core 2 *O*-glycans in relation to other shorter glycans, thus like stressed cell lines (**Fig. 6h; Supplementary Table 8**). The major difference was the absence of core 3 in tissues, abundant in cell lines, and dominant sialyl-T antigens, denoting a tissue specific regulation. Trace amounts of Tn and STn were also detected but no fucosylated T antigens, as previously found *in vitro*. Contrastingly, HIF-1α^negative^/Ki-67^high^ tumours were more heterogeneous, with some tumours showing hypoxic-like glycopatterns while others exhibited high percentage of core 2 elongations, suggesting that other l factors may also drive altered glycosylation. Still, immunohistochemistry screening of tissue sections showed an increase in Tn as well as sialylated T expressions in hypoxic tumours in comparison to normoxic, accompanied by the loss of core 2 and terminal Lewis antigens, which were used as surrogates of extended *O*-glycans (**Figs. 6j and k**). In agreement with these observations, double immunofluorescence confirmed higher levels of Tn and sialylated T antigens in hypoxic tumour areas after *N*-deglycosylation with PnGase F (**Fig. 6l**). In addition, we performed a broad characterization of the *N*-glycome. Despite significant structural diversity, hypoxic tumours were enriched for oligomannose *N*-glycans (**Fig. 6i; Supplementary Table 9**), which was not evident *in vitro*. Collectively, these observations highlight fundamental microenvironment-driven molecular grounds shared by cell models and tumours, including the adoption of an immature *O*-glycocalyx and potentially N-glycosylation, which warrants deeper investigation.

### Stress-induced glycome alterations contribute to cancer aggressiveness

A library of T24 cells (**Supplementary Figs. 8 and 9**) displaying distinct simple *O*- glycophenotypes was constructed to gain more knowledge on the biological role played by altered glycosylation in BLCA. Notably, we also tried to edit 5637 cells without success. We started by overexpressing *ST3GAL1* (**Supplementary Fig. 10**), which decreased the T antigen and core 1 fucosylation but not core 2-derived glycans, thus not reflecting the hypoxic phenotype (**Supplementary Table 10**). Therefore, we glycoengineered cells to hamper *O*-glycan extension beyond core 1 and core 2 by supressing *C1GALT1* (**Supplementary Fig. 8**) and *GCNT1* (**Supplementary Fig. 9**), respectively, using validated gRNAs^48^ through CRISPR-Cas9 technology. T24 *C1GALT1* KO cells were invariably characterized by a marked increase in Tn antigen, with minor changes in STn and core 3 expressions (**Fig. 7a**). Core 3 and STn antigens emerge as the dominant ions in glycome analysis, reflecting the fact that the Tn antigen cannot be detected using our methodology. Also, as expected, further extension to core 1 was not observed **(Figs. 7a and b)**. *GCNT1* KOs resulted in the conservation of sialylated T antigens (**Figs. 7i and j; Supplementary Table 10**), with similar percentages of mono and di-sialylated species as observed in hypoxia, and complete loss of core 2 and other extended glycans. High levels of fucosylated core 1 could still be detected. Interestingly, *ST3GAL1* overexpression on T24 wild type cells decreased core 1 and fucosyl- core 1, increasing the percentage of sialylated core 2 glycans, whereas for *GCNT1* KO cells it increased T antigen di-sialylation (**Supplementary Table 10).** These observations strongly suggest that *ST3GAL1* elevation alone cannot account for the stress-associated glycome.

**Fig. 7.**
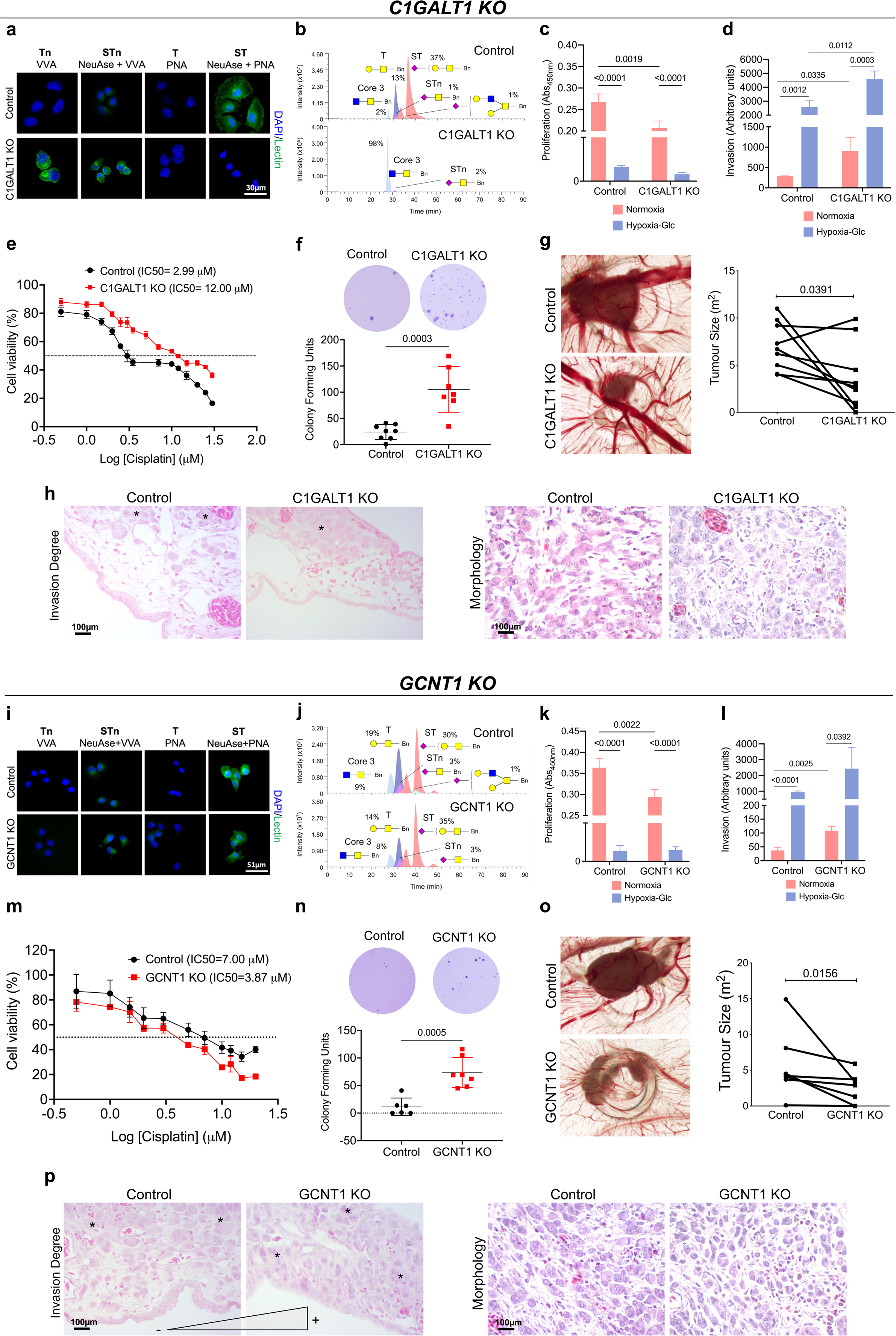
Hypoxia and low-glucose-induced simple glycophenotypes drive relevant cancer-associated hallmarks. a, b. T24 *C1GALT1* KOs show complete abrogation of *O*- glycans extension beyond the Tn antigen, mirroring a major alteration observed in hypoxic tumours. Glycoengineered cells homogeneously express the Tn antigen and show low levels of core 3. Mock controls’ glycosylation closely resembles T24 wild-type cells. **c.** *C1GALT1* KOs display reduced proliferation compared to controls under normoxia. Collectively, altered glycosylation impacts more in proliferation when compared to hypoxia and glucose deprivation. **d.** Under normoxia and hypoxia with glucose deprivation, *C1GALT1* KOs glycoengineered cells display significantly enhanced invasion, suggesting a critical role of *C1GALT1* in modulating invasive behaviour under stress. **e, f.** *C1GALT1* KOs demonstrate higher resistance to cisplatin (**e**) and anoikis compared to mock controls (**f**). **g,h.** In CAMs, *C1GALT1* KOs give rise to smaller tumours (**g**), in agreement with proliferation studies *in vitro* (**c**), showing less cohesive features and invasive patterns compared to control (**h**). **i, j.** T24 *GCNT1* KOs exhibit complete abrogation of *O*-glycans extension beyond core 1, mirroring another major alteration observed in hypoxic tumours. As a result, glycoengineered cells express high levels of sialylated T antigens, namely sialyl-T, but do not present core 2-derived glycans. Mock controls’ glycosylation closely resembles T24 wild-type cells. **k, l.** *GCNT1* KOs display reduced proliferation compared to controls (**k**) and increased invasion (**l**) under normoxia, resembling *C1GALT1* KOs. A higher invasion is also observed under hypoxia -Glc. **m, n.** *GCNT1* KOs demonstrate lower resistance to cisplatin (**m**) but higher resistance to anoikis compared to mock controls (**n**). **o, p.** In CAMs, *GCNT1* KOs give rise to smaller (**o**), and more invasive tumours compared to controls (**p**). Unpaired T-test, Two-way ANOVA followed by Tukey’s multiple comparison test, and Wilcoxon test were used for statistical analysis.

*C1GALT1* and *GCNT1* KO cells closely mimicking the glycophenotype of hypoxic cells were further used to interrogate the role played by glycosylation in decisive aspects of the disease, namely capacity to proliferate, invade, grow without anchorage, potentially metastasise, and tolerate chemotherapy agents (**Fig. 7**). Both were less proliferative *in vitro* under normoxia as well as hypoxia and low glucose (**Figs. 7c and k)**. Higher capacity to grow in an anchorage-independent manner and resist anoikis was also observed, supporting increased metastatic potential **(Figs. 7f and n)**. *C1GALT1* KO cells were more resistant to cisplatin over a wide range of drug concentrations and presented higher IC50 than its control **(Fig. 7e)**. Increased invasion was also observed under normoxia and microenvironmental stress (**Fig. 7d**). Complementary phosphoproteome characterization confirmed enrichment for pathways consistent with these functional alterations (**Supplementary Fig. 11a**), associated to activation of relevant tyrosine kinases also found in hypoxic cells and tumours (**Supplementary Fig. 11b**), suggesting common molecular grounds supported by immature glycosylation. Interestingly, *GCNT1* KO also presented low proliferation and higher invasion capacity in normoxia and hypoxia plus low glucose(**Fig. 7l**), but not increased resistance to cisplatin **(Fig. 7m)**. However, the molecular pathways governing glycan-linked functional alterations differ significantly from those in Tn overexpressing cells (**Supplementary Fig, 11c-d**), thus emphasizing the crucial role of glycosylation in fine-tuning molecular functions.

Finally, we explored the *in ovo* chick chorioallantoic membrane (CAM) assay to assess the biological role of glycans *in vivo*. Notably, T24 CAM models have been previously developed and proven to reflect many histological and molecular features of human bladder tumours^49^, providing relevant platforms for functional studies. *C1GALT1* and *GCNT1* KO CAM models exhibited significantly smaller tumours when compared to controls, which was consistent with observations *in vitro* **(Figs. 7g and o)**. Interestingly, despite display much smaller lesions, *C1GALT1* KO tumours invaded similarly to controls and exhibited a less cohesive phenotype characterized by a higher number of isolated tumour niches, suggesting higher motility (**Fig. 7h**). *GCNT1* KO models invaded more, reaching the lower allantoid epithelium (endoderm) and, in some cases, even expanding beyond that **(Fig. 7p**). In summary, we have highlighted the different contributions of distinct glycosylation patterns associated with hypoxia and glucose suppression to cancer progression and potentially dissemination. The induction of short glycosylation also had a major impact on cell proliferation as potentiated invasion, in perfect agreement with the observation of these glycans in hypoxic tumour areas. These cells also showed higher metastatic potential as well as higher tolerance to cisplatin, in agreement with observations made for similar models from other solid tumours^50,51^.

## Discussion

Hypoxia and glucose deprivation are relevant features of solid tumours, arising from sustained proliferative signalling and defective neo-angiogenesis, which decisively shape the molecular and functional phenotypes of tumour cells. The contribution of hypoxia to disease severity, even though extensively studied and rather well known, has been addressed without considering the concomitant influence of low glucose. Acknowledging the complex microenvironmental challenges experienced by cells striving to survive in poorly vascularized tumour areas, the present study devoted to understanding cells’ plasticity and adaptability to these selective pressures, providing means for more educated and precise interventions. We have comprehensively addressed the impact of microenvironmental stressors (hypoxia and low glucose) at the transcriptome, metabolome and phosphoproteome levels, which ultimately led us to interrogate the glycome and its biological implications in disease. Strikingly, BLCA models of distinct genetic and molecular backgrounds responded similarly to microenvironmental challenge, denoting common biological grounds facing stress. Cells tolerated low levels of oxygen and glucose, maintaining viability, and avoiding apoptosis, while dramatically decreasing proliferation. Concomitantly, cells became more invasive, denoting an active strategy to escape suboptimal growth conditions. Stress also increased capacity to tolerate the chemotherapy drug cisplatin, possibly explaining the resilient nature of poorly vascularized bladder tumours^52,53^. Interestingly, this aggressive behaviour was completely reverted by re-oxygenation and glucose restoration without compromising cell viability. This further highlighted relevant molecular plasticity accommodating microenvironmental challenges. Under stress, despite the low levels of oxygen, cancer cells adopted mitochondrial fatty acid *β*-oxidation and oxidative phosphorylation rather than anaerobic glycolysis as main bioenergetic pathway, explaining the tremendous decrease in lactate production by these cells. Similar observations were made for prostate cancer, where fatty acid oxidation is a dominant bioenergetic pathway rather than glycolysis, showing potential to be the basis of imaging diagnosis and targeted treatment^54^. To some extent, this also challenges the pivotal role attributed to lactate in the aggressiveness of hypoxic cancer cells in many studies^55^. We have also found strong indications of mitochondrial autophagy, which has been previously described as a survival mechanism of cancer cells under nutrient deprivation, driving invasion, immune escape, and metastasis^11^. Notably, most of the stress-induced molecular alterations found *in vitro* were observed in highly hypoxic and aggressive bladder tumours, providing a basis for patient stratification and targeted interventions.

Finally, we have shown that both stressed BLCA and hypoxic tumours exhibit immature glycophenotypes, characterized by the incapacity to extend *O*-glycans beyond core 2. In previous studies, we have linked short glycoforms to advanced-stages, which are frequently associated with poor prognosis^13,56–58^. However, the microenvironmental triggers for glycocalyx remodelling remained to be elucidated. Here, we demonstrate that hypoxia and glucose deprivation are key drivers of these alterations by promoting changes in glycogenes expression and limiting nucleotide sugar donors due to low glucose levels, potentially aggravated by a disorganization of secretory pathways. Changes in protein *O*-glycosylation may be part of a wide array of responses that make BLCA cells more capable of resisting microenvironmental stress. Specifically, these enabled cells to halt proliferation, resist anoikis, and tolerate chemotherapy. Shorter *O*-glycosylation also provides a stress-escape mechanism by promoting less cohesive and invasive traits. These observations are in perfect agreement with other reports from simple cell cancer models of different origins, highlighting their pancarcinomic nature^59^. These are seminal findings that offer new insights into understanding the drivers of cancer-associated glycosylation, which have so far been linked to functional mutations in *COSMC*^60^ and, more recently, pro-inflammatory immune responses^61^. Additionally, the Tn antigen found in hypoxic cells is considered a key mechanism of cancer immunosuppression, impairing the maturation of macrophages and dendritic cells by interacting with immune system lectins^62^. Future studies on the hypoxia-glycome-immune response axis will likely offer novel insights into the capacity of cells to thrive under stress conditions. In summary, this study provides the microenvironmental context for previous observations regarding mucin-type *O*-glycan expression in bladder tumours, with potential implications for other solid tumours where short-chain glycans also play key functional roles. Furthermore, it lays the groundwork to identify quiescent cells in hypoxic niches, which may eventually be responsible for disease recurrence upon reoxygenation, paving the way for novel targeted interventions.

## Acknowledgments

The authors wish to acknowledge the Portuguese Foundation for Science and Technology (FCT) for PhD grants SFRH/BD/111242/2015 (AP), 2020.09384.BD (DF), 2022.12980.BD (AM), SFRH/BD/146500/2019 (MRS), 2020.08708.BD (RF), SFRH/BD/127327/2016 (CG), 2020.09394.BD (SC), SFRH/BD/142479/2018 (JS), Principal researcher contract 2022.08311.CEECIND (JAF), and junior researcher contact 2021.03835.CEECIND (AB). AP acknowledges the CI-IPOP junior researcher position UIDP/00776/2020–5. FCT is co-financed by European Social Fund under Human Potential Operation Programme from National Strategic Reference Framework. The authors also acknowledge FCT/MCTES funding within the projects RESOLVE (PTDC/MED-OUT/2512/2021) and REVERENT (2022.03621.PTDC) and funding for the IPO research center (PEst-OE/SAU/UI0776/201, CI-IPOP-29-2016-2022, CI-IPOP-58- 2016-2022), for CQUM (UID/QUI/00686/2020), and the LAQV research unit (UIDB/50006/2020 | UIDP/50006/2020). We further acknowledge funding for the large-scale proteomics facility, the Netherlands Proteomics Center, through the EU Horizon 2020 program Epic-XS (project 823839). This article is also a result of the project NORTE-01-0145-FEDER-000012 and “The Porto Comprehensive Cancer Center Raquel Seruca” with the reference NORTE-01-0145-FEDER- 072678 - Consórcio PORTO.CCC—Porto.Comprehensive Cancer Center Raquel Seruca, supported by NORTE 2020, under the PORTUGAL 2020 Partnership Agreement, through the European Regional Development Fund. This work was also financed by FEDER through the COMPETE 2020 - Operational Programme for Competitiveness and Internationalisation (POCI- 01-0145-FEDER-007274 for i3S unit). Finally, the authors also acknowledge the ICBAS PhD Program in Biomedical Sciences and the support of the i3S scientific platforms for the CAM assays and HEMS (PPBI-POCI-01-0145-FEDER-022122). Moreover, the authors acknowledge the kind collaboration of the pharmaceutical units of IPO-Porto for providing and supervising the handling of clinical grade cisplatin.

## Author Contributions

JAF conceived and supervised the study; AP, DF, MRS, AMNS performed glycomics analysis, AP, DF, AM, CP, EF, SC, RF, and CG performed immunoassays; AP, MRS, TSV, and AJRH conducted phosphoproteomics analysis; MC, and PP aided AP with genetic editing; AP, DF, AM, AB, MRS, JS, LL, FT, JAF and AMNS curated the data, performed bioinformatics, and statistical analysis; RF and MJO gave valuable scientific input and provided technological resources for project execution; LLS and JAF gathered the financial and human resources for the study; AP, AM, and JAF wrote the manuscript, which was reviewed and commented by all the co-authors.

## Methods

### Cell culture

Human BLCA cell lines RT4, 5637, T24, and HT1197 were purchased from American Type Culture Collection (ATCC). RT4, 5637, and T24 cells were maintained with complete RPMI 1640 GlutaMAX™ medium (Gibco) and HT1197 with DMEM GlutaMAX™ medium (Gibco) supplemented with 10% FBS (Gibco). Cells were kept at 37 °C in a 5% CO_2_ humidified atmosphere (Normoxia). Cells were also grown for 24h under hypoxia and nutrient deprivation (Hypoxia -Glc) at 37 °C in a 5% CO_2_, 99.9% N_2_ and 0.1% O_2_ atmosphere, using a BINDER C- 150 incubator (BINDER GmbH), and complete RPMI 1640 or DMEM media without glucose (Gibco). A 24h exposure to 500 µM deferoxamine mesylate salt (DFX, Sigma) in complete medium was used as positive control for HIF-1α stabilization. For re-oxygenation experiments, cells under oxygen and glucose deprivation were restored to standard culture conditions 24 h prior to analysis.

### HIF-1α expression

HIF-1α was evaluated using a HIF-1 Alpha ELISA Kit (Invitrogen™). Procedure steps were followed according to the manufacturer’s instructions and results were monitored at 450 nm. The results were normalized to cell proliferation rates.

### Proliferation assay

Cell proliferation was determined using a colorimetric Cell Proliferation ELISA (Roche) kit. Procedure steps were followed according to the manufacturer instructions and results were monitored at 450 nm.

### Lactate Assay

A colorimetric Assay Kit (Abcam) was used to detect L(+)-Lactate in deproteinized cultured cells lysates and conditioned medium. Procedure steps were followed according to the manufacturer’s instructions and results were monitored at 450 nm. The results were normalized to cell proliferation rates.

### Invasion Assay

Invasion assays were performed using Corning® BioCoat™ Matrigel® Invasion Chambers as described in Peixoto, A. *et al*.^13^. Invasion counts were normalized to cell proliferation rates, and cells were seeded in quintuplicates for each experiment.

### Anchorage-independent growth

Anchorage-independent growth was measured using the soft agar colony formation assay. A 0.5% low melting point agarose (Lonza) solution in complete medium was used as bottom layer in 6- well flat bottom plates. A top layer of 0.3% agarose containing 1x10^4^ cells was then plated and covered with culture medium. Cells were maintained in standard growth conditions for one month. Colonies were then fixed with 10% neutral buffered formalin solution (Sigma-Aldrich) and stained with 0.01% (w/v) crystal violet (Sigma-Aldrich). Colonies were photographed using a stereomicroscope (Olympus, SZX16 coupled with a DP71 camera) and automatically counted using the open-source software ImageJ (Fiji package). Only colonies containing more than 50 cells were considered.

### Cell viability assay

Cell viability was determined using the Trypan Blue Exclusion Test of Cell Viability. For specific apoptosis stage assessment, cells were detached using Accutase enzyme cell detachment medium (Thermo Fisher Scientific) and apoptosis was determined using the Cell Apoptosis Kit with FITC annexin V and propidium iodide for flow cytometry (Thermo Fisher Scientific), according to the manufacturer’s instructions. Data analysis was performed through CXP Software in a FC500 Beckman Coulter flow cytometer.

### Cisplatin resistance

BLCA cells were plated into 96-well plates, following a 24h exposure to crescent concentrations of cisplatin. Positive and negative controls of cell death were set, consisting of cell incubation with complete medium with and without 1% Triton-X (Sigma-Aldrich), respectively. After cisplatin incubation, cells were incubated with 1.2mM 3-(4,5-dimethylthiazol-2-yl)-2,5- diphenyltetrazolium bromide (MTT, Thermo Fisher Scientific) solution. Formazan crystals were solubilized with dimethyl sulfoxide (DMSO, Sigma-Aldrich), following absorbance measurement at 540 nm. The percentage of cell viability was calculated as follows:

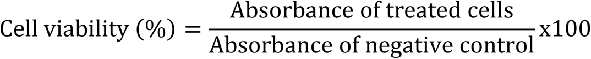

### Transcriptomics

Transcriptomics study was performed according to the protocol previously described by us^63^.

### Metabolomics

Cells were dispersed in 80% methanol (Merck), sonicated for 30 min at 4 °C and kept at -20 °C for 1 h. Samples were then centrifuged, and the supernatant was analysed by UHPLC-ESI-MS/MS in positive and negative mode. Metabolite analysis was performed using an Ultimate 3000 LC combined with QExactive mass spectrometer (Thermo Fisher Scientific). Eluent A was 0.1% formic acid in water and eluent B was acetonitrile. Metabolite separation occurred using the following gradient elution (0-1.5 min, 95-70% A; 1.5-9.5 min, 70-5% A; 9.5-14.5 min, 5% A; 14.5-14.6 min, 5-95% A; 14.6-18.0 min, 95% A). The flow rate of the mobile phase was 0.3 mL/min. The column (Acquity UPLC HSS T3; 100 Å, 1.8 μm, 2.1 mm × 150 mm) temperature was maintained at 40 °C, and the sample manager temperature was set at 4 °C. Metabolites were identified by retention time and corresponding MS/MS spectra. For metabolomics data pre- processing and analysis, raw data matrices were blank subtracted (a mean blank value was calculated per metabolite) and normalized to the number of cells for each condition. The resulting matrices were then imported to Metaboanalyst 4.0 and log-transformed to reduce heteroscedasticity and pareto-scaled to adjust for differences in fold-changes between metabolites.

### Metabolome and Metabolome/Transcriptome joint pathway analysis

Multivariate and univariate analysis were performed to identify metabolites that discriminate normoxia from hypoxia with low glucose. Unsupervised principal components analysis (PCA) was applied to unravel data structure, following a supervised method, namely partial-least-squares discriminant analysis (PLS-DA) to identify which metabolites were useful to predict group membership. Metabolites with discriminative power were ranked based on Variable Importance in Projection values >1 and PLS-DA models were validated based on the “prediction accuracy during training” test statistic with 1000 permutations (*p*<0.05 for significance). Heat maps with hierarchical clustering of metabolites were constructed based on: i) distance measure: Pearson correlation (similarity of expression profiles), ii) clustering algorithm: complete linkage (forms compact clusters), iii) feature autoscale. Hierarchical clustering of samples was carried out based on: i) distance measured: Euclidean distance (sensitive to magnitude differences), ii) clustering algorithm: Ward (minimizes within-cluster variance). Differences in metabolites between groups were further evaluated using ne-way analysis of variance (ANOVA) with a False Discovery Rate (FDR) cut-off set at 0.05 for significance. Tukey’s post hoc was applied to check which groups differed. Significant metabolites unravelled by ANOVA were then used for pathway analysis to identify the most relevant pathways that are involved in the adaptation of cells from normoxia to hypoxia with low glucose. Pathway analysis was carried out based on: i) functional enrichment, which was assessed using hypergeometric test for over-representation analysis (*p*<0.05 for significance) and ii) pathway topology analysis, which was implemented using the relative betweenness centrality. Pathway impact was considered relevant if >0.1. The joint pathway analysis was carried out using transcriptomic and metabolomic data, based on a gene and metabolite list with associated fold-changes. The human pathway library was chosen using the pathway database “all pathways (integrated)”. The enrichment analysis was based on the hypergeometric statistic test, while degree centrality was used as topology measure. The integration method was based on a combination of queries.

### ATP quantification assay

A fluorometric ATP assay kit (Abcam) was used according to the manufacturer’s instructions to determine ATP levels in deproteinized whole cell lysates. Fluorimetric products were quantified using a microplate reader. The results were normalized to cell proliferation.

### Western Blot

Whole protein extracts were collected from BLCA cells using a 25 mM Tris-HCl pH 7.2, 150 mM NaCl, 5 mM MgCl_2_, 1% NP-40, and 5% glycerol lysis buffer, supplemented with Halt™ Protease and Phosphatase Inhibitor Cocktail (Thermo Fisher Scientific). Isolated proteins were run on 4– 20% precast SDS-PAGE gels (Bio-Rad) and screened for AMPK, phosphorylated AMPK, and a panel of glycosyltransferases (C1GalT1, Cosmc, B3GnT6, C2GnT, ST3Gal-I). The antibodies and experimental conditions are detailed in **Supplementary Table 11**. Total protein stain was assessed using Ponceau S anionic dye.

### Citrate synthase assay

Citrate synthase activity was measured in whole cell protein lysates using the method proposed by Coore *et al.*^64^. In brief, the coenzyme A released from the reaction of acetyl-CoA (Sigma-Aldrich) with oxaloacetate (Sigma-Aldrich) was determined by its reaction with 5,5’-dithiobis-(2- nitrobenzoic acid) (Sigma-Aldrich) at 412 nm (ε=13.6 mM^-1^cm^-1^). The results were normalized to cell proliferation.

### Transmission Electron Microscopy

For electron microscopy, T24 and 5637 cells were fixed in 2% glutaraldehyde (Electron Microscopy Sciences) with 2.5% formaldehyde (Electron Microscopy Sciences) in 0.1 M sodium cacodylate buffer (pH 7.4). Cells were post-fixed in 1% osmium tetroxide (Electron Microscopy Sciences) diluted in 0.1 M sodium cacodylate buffer. Samples were then dehydrated and embedded in Epon resin (TAAB). Ultra-thin 50 nm sections were cut on an RMC Ultramicrotome (PowerTome) using Diatome diamond knives, mounted on 200-mesh copper grids (Electron Microscopy Sciences), and stained with uranyl acetate substitute (Electron Microscopy Sciences) and lead citrate (Electron Microscopy Sciences). Sections were then examined under a JEOL JEM 1400 transmission electron microscope (JEOL), and images were digitally recorded using a CCD digital camera Orius 1100W.

### Phosphoproteomics and data analysis

The preparation of samples for phosphoproteomics was performed according to Veth, T. S. *et al.*^65^. Then, samples were suspended in 2% formic acid and analysed on an Exploris (Thermo Scientific) coupled to an UltiMate 3000 (Thermo Scientific), fitted with a µ-precolumn (C18 PepMap100, 5µm, 100 Å, 5mm × 300µm; Thermo Scientific), and an analytical column (120 EC-C18, 2.7µm, 50cm × 75µm; Agilent Poroshell). Peptides were loaded in 9% Buffer A (0.1% FA) for 1 min and separated using 9-36% buffer B (80%ACN, 0.1%FA) in 97 min at 300 nL/min, followed by a 6 min column wash with 99% buffer B at 300 nL/min, and a 10-minute column equilibration at 9% B. The MS was operated in DDA mode, with the MS1 scans in a range of 375-1600 m/z acquired at 60k, using an automatically set AGC target. MS2 scans were acquired with a 16s dynamic exclusion at a 30k resolution, 28% normalized collision energy, and an isolation window of 1.4 m/z. Raw files were processed via MaxQuant version 1.6.17.0 using the verified human proteome from UniprotKB (release 09-2019) containing 20,369 proteins^66^. A maximum of 5 modifications and two missed cleavages were set, using fixed carbamidomethyl modification, and the variable modifications oxidized methionine, protein *N*-terminal acetylation, and serine/threonine/tyrosine phosphorylation. The protein and peptide FDR were set to <0.01 and conducted with match between runs enabled. Using the Perseus framework, only class 1 phosphosites were included (>0.75), and with minimally two detections in at least one group for subsequent analysis. Missing values were replaced using the normal distribution and a downshift of 1.8. The data was subsequently explored using Phosphomatics^67^ and xxx

### Glycomics

The *O*-glycome of BLCA cells challenged by microenvironmental stressors as well as glycoengineered cell models was characterized through the Cellular *O*-glycome Reporter/Amplification method^68^, using the protocol detailed in Fernandes, E. *et al.*^69^. The *O*- glycome of oesophageal, gastric, and colorectal cancer cells as well as human monocyte-derived macrophages was also characterized following the same protocol. Regarding bladder tumours, *N*- and *O*-glycomics was performed as previously described^70^. Briefly, FFPE tissues were deparaffinized, rehydrated, and subjected to heat-induced antigen retrieval using a citrate-based solution (Vector Laboratories). Then, proteins were denatured and reduced by incubation with 150 µl denaturation mix (145 µl of 8 M GuHCl and 5 µl of 200 Mm DTT) at 60 °C for 30 min. *N*- glycan release was achieved after digestion with PNGase F (1 U/10 µg protein at 37 °C overnight; Promega). Released *N*-glycans were hydrolysed with 25 µl of 100 mM ammonium acetate at pH 5 (1 h at RT), removing the glycosylamine form of the non-reducing terminus, and subsequently reduced with 20 µl of 1 M NaBH_4_ in 50 mM KOH (3 h at 50 °C). The reaction was quenched by adding glacial acetic acid. *O*-glycans were released by reductive β-elimination (1 M NaBH_4_ in 50 mM KOH overnight at 50 °C). Finally, reduced *N*- and *O*-glycans samples were desalted using a cation exchange resin (AG 50W-X8; Bio-Rad). *N*-glycomics of BLCA cell lines was performed similarly after immobilization of membrane protein extracts in a hydrophobic PVDF membrane (MAIPS4510; Millipore). All glycans were permethylated and analysed by reverse phase nanoLC-ESI-MS/MS as previously described by us^69^, using a 3000 Ultimate nano-LC coupled to either an LTQ-Orbitrap XL or QExactive mass spectrometer (Thermo Fisher Scientific). Glycans structures were identified considering previous knowledge on glycosylation, chromatography retention times, *m/z* identification, and corresponding product ion spectra. The relative abundance resulted from the sum of the extracted ion chromatogram areas for each glycan structure in relation to the sum of chromatographic areas of all identified glycans.

### Flow cytometry for glycans detection

BLCA cells were screened for Tn, STn, T, and ST antigens as well as for *N*-acetylglucosamine residues by flow cytometry as previously described in Peixoto, A *et al.*^63^. The antibodies and experimental conditions are detailed in **Supplementary Table 11**. For evaluation of STn expression, an isotype control and cells digested with neuraminidase prior to STn staining were used as negative controls. Regarding GSL II lectin detection, PNGaseF enzymatic digestion was performed to exclude *N*-glycan-associated *N*-acetylglucosamine residues contribution. Data analysis was performed through CXP Software in a FC500 Beckman Coulter flow cytometer.

### Immunocytochemistry and histochemistry

Immunofluorescence (IF) was performed to evaluate Tn, STn, T and sialyl-T antigens in T24 glycoengineered cell models using the same lectins and strategies employed in flow cytometry. Cells were cultured at low density and fixed with 4% paraformaldehyde (Sigma-Aldrich). After antigen staining, cells were marked with 2,3x10^-3^ µg/µL 4’,6-Diamidino-2-phenylindole dihydrochloride (DAPI, Thermo Fisher Scientific) for 10 min at RT in the dark. Sialylated glycoforms were evaluated in parallel with samples digested with 50 mU/mL neuraminidase overnight at 37 °C. All images were acquired on a Leica DMI6000 FFW microscope using Las X software (Leica). Furthermore, BLCA FFPE tissue sections were screened for Tn, STn, T, ST, SLe^A^, SLe^X^, HIF-1α, and KI-67 antigens by immunohistochemistry (IHC) using a previously described streptavidin/biotin peroxidase method^58^. Tumours were also screened by immunofluorescence envisaging Tn, ST, and HIF-1α co-detection. Immuno-reactive sections were blindly assessed using light microscopy by two independent observers and validated by an experienced pathologist. Antibodies and experimental conditions used for immunoassays are detailed in **Supplementary Table 11**.

### RT-PCR for glycogenes expression

*C1GALT1*, *C1GALT1C1*, *B3GNT6*, *GCNT1*, *GCNT2*, *GCNT3*, *GCNT4*, *B3GNT3*, *FUT1*, *FUT2*, *FUT3*, *FUT4*, *FUT5*, *FUT6*, *FUT7*, *FUT9*, *ST3GAL1*, *ST3GAL3*, *ST3GAL4*, *ST6GANAC1*, *ST6GALNAC2*, *ST6GALNAC3*, and *ST6GALNAC4* gene expression was assessed by quantitative polymerase chain reaction. Briefly, total RNA from cultured cells was isolated using TriPure isolation Reagent (Roche), according to the manufacturer’s instructions. RNA conversion and gene expression analysis were performed using TaqMan™ Gene Expression Assays (Applied Biosystems™). The mRNA levels were normalized to the expressions of *B2M* and *HPRT* housekeeping genes. Taqman probes used for the glycogenes screening are detailed in **Supplementary Table 12**.

### Patient Samples and Ethics

A retrospective series of 70 formalin-fixed paraffin-embedded bladder tumours (FFPE; 30 NMIBC; 40 MIBC) from the IPO-Porto biobank were used for this study after patient’s informed consent (ethics project reference: CES 86/017).

### Chicken embryo chorioallantoic membrane (CAM) assay

The CAM assay was used to assess the *in vivo* establishment of tumour aggregates derived from glycoengineered cell models. On embryonic development day (EDD) 3, a squared window was opened in the eggshell of fertilized chicken (*gallus gallus*) eggs, and 2–2.5 mL of albumen was removed to allow detachment of the developing CAM. The window was then sealed with adhesive tape and the eggs were incubated horizontally at 37.8 °C in a humidified atmosphere. On EDD9, 1x10^6^ cells derived from each developed cell model were re-suspended in 10μl of Corning® Matrigel® Matrix and placed in a 3 mm silicone ring attached to the growing CAM. Control cells and respective glycoengineered models were inoculated in the same egg. At least 10 viable embryos were used per experimental pair. The eggs were re-sealed and returned to the incubator for one week. On EDD16, the CAM was excised from the embryos, photographed *ex-ovo* under a stereoscope at 20x magnification (Olympus, SZX16 coupled with a DP71 camera), and images were analysed to determine tumour size. CAM attached tumours were then formalin fixed and paraffin embedded and stained with haematoxylin and eosin.

### Public Gene Expression and Clinical Data Statistical Analysis

Normalized gene expression data and the corresponding clinical data, including survival information, for 407 bladder cancer patients were obtained from The Cancer Genome Atlas (TCGA) database using the Xena browser. In addition, normalized gene expression data from healthy bladder tissue samples (n=9) from the Genotype-Tissue Expression (GTEx) project was also retrieved. Univariable Cox regression analysis was used to analyze and identify a survival-associated hypoxia gene signature. Hazard ratios (HRs) and corresponding 95% confidence intervals (CIs) were estimated for the 30 most differentially expressed genes in hypoxia and low glucose conditions using the “finalfit” R package. Kaplan-Meier (KM) analysis and log-rank test were performed to evaluate the survival outcomes in the patients with the hypoxia gene signature identified. The optimal cutoffs to categorize the expression of each gene in “high” or “low” were determined by the “surv_cutpoint” algorithm of the “survival” R package. Heatmaps were constructed using the “ComplexHeatmap” R package, to assess gene expression profile between conditions, hypoxia versus normoxia. *p*-value <0.05 was considered statistically significant. All statistical analyses were performed using R (version 4.2.1).

### Statistical Analysis

Statistical analyses were performed using the GraphPad Prism software. One-way ANOVA followed by Tukey’s multiple comparisons test was used in evaluation of HIF-1α, lactate, proliferation, invasion, and the autophagy experiment. The Mann-Whitney test was used in the analysis of cisplatin IC50, L-carnitine and citrate levels, glycosyltransferases expression, AMP/ATP ratio, glycans expression data from flow cytometry assay, and relative abundance of *O*- and *N*-glycans extension in tumours. The Mann-Whitney test was also used in the evaluation of AMPK expression and the Two-way ANOVA followed by Tukey’s multiple comparisons test was used for glycoengineered cells’ proliferation and invasion. Finally, unpaired t test was used in the analysis of percentage of HIF-1α-positive tumour area and for colony forming assays. CAM assay results were analysed using a Wilcoxon test. Statistical analysis of phosphoproteomics data was performed in Phosphomatics software^67^. Differences were considered significant for *p*<0.05. All experiments were performed in triplicates and three replicates were conducted for each independent experiment. The results are presented as the average and standard deviation of these independent assays.

## Supplementary information

Supplementary Data and Supplementary Methods (cell cycle analysis; autophagy assays, and BLCA cells glycoengineering) corresponding to experiments’ results shown in supplementary information are available for this paper.

## Data availability (to be updated)

Transcriptomics data is under submission to GEO (https://www.ncbi.nlm.nih.gov/geo/). Metabolomics data is under submission to Metabolomics Workbench (https://www.metabolomicsworkbench.org/). Glycomics is under submission to GlycoPost (https://glycopost.glycosmos.org/). Phosphoproteomics data is available in PRIDE (accession: PXD045777).

