## Supplemental Methods for "Multilevel Plasticity and Altered Glycosylation Drive Aggressiveness in Hypoxic and Glucose-Deprived Bladder Cancer Cells"

### Supplementary Methods

Andreia Peixoto[1,2,3], Dylan Ferreira,[1,2,3#], Andreia Miranda[1,2,3#], Marta Relvas-Santos[1,2,3,4], Rui Freitas[1,2,3], Tim S. Veth[5,6], Andreia Brandão[1], Eduardo Ferreira[1], Paula Paulo[1], Marta Cardoso[1], Cristiana Gaiteiro[1,3], Sofia Cotton[1,3], Janine Soares[1,3,7], Luís Lima[1], Filipe Teixeira[8], Rita Ferreira[7], Carlos Palmeira[1,9,10], Albert J. R. Heck[5,6], Maria José Oliveira[2], André M. N. Silva[4], Lúcio Lara Santos[1,10,11], José Alexandre Ferreira[1,3*]

[1]Research Center of IPO-Porto (CI-IPOP) / RISE@CI-IPOP (Health Research Network), Portuguese Oncology Institute of Porto (IPO-Porto) / Porto Comprehensive Cancer Center (P.ccc) Raquel Seruca, Porto, Portugal; [2]i3S – Instituto de Investigação e Inovação em Saúde, Universidade do Porto, Porto, Portugal; [3]Institute of Biomedical Sciences Abel Salazar (ICBAS), University of Porto, Porto, Portugal; [4]LAQV-REQUIMTE, Department of Chemistry and Biochemistry, Faculty of Sciences, University of Porto, Porto, Portugal; [5]Biomolecular Mass Spectrometry and Proteomics, Bijvoet Center for Biomolecular Research and Utrecht Institute for Pharmaceutical Sciences, University of Utrecht, Utrecht, The Netherlands; [6]Netherlands Proteomics Center, Padualaan, Utrecht, The Netherlands; [7]QOPNA & LAQV-REQUIMTE, Department of Chemistry, University of Aveiro, Aveiro, Portugal; [8]Centre of Chemistry, University of Minho, Braga, Portugal; [9]Department of Immunology, Portuguese Oncology Institute of Porto, Porto, Portugal; [10]Health School of University Fernando Pessoa, Porto, Portugal; [11]Department of Surgical Oncology, Portuguese Oncology Institute of Porto, Porto, Portugal.

#equal contribution

*corresponding author

#### Cell cycle analysis

Cells were harvested by trypsinization, following fixation and labelling with DNA labelling solution (Cytognos) for 10 min at RT in obscurity. Data analysis was performed through CXP Software in a FC500 Beckman Coulter flow cytometer.

**Glycoengineered cell models**

A recombinant *Streptococcus pyogenes* Cas9 (GeneArt™ Platinum Cas9 Nuclease, Thermo Fisher Scientific) and single-guided RNA (GTAAAGCAGGGCTACATGAG, sgRNA) were used to generate site-specific double-strand breaks in the C1GALT1 gene in T24 cells *in vitro*. In parallel, two sgRNAs were used for GCNT1 gene knock-out (KO) (gRNA1: TAGTCGTCAGGTGTCCACCG, gRNA2: AAGCGGTATGAGGTCGTTAA). Lipofectamine™ CRISPRMAX™ Transfection Reagent (Thermo Fisher Scientific) was used according to the manufacturer's instructions. Complexes were made in serum-free media (Opti-MEM™ I Reduced Serum Medium) and added directly to cells in culture medium. Single clones were obtained by serial dilution in 96-well plates and KO clones were identified by Indel Detection by Amplicon Analysis (IDAA) using ABI PRISM™ 3010 Genetic Analyzer (Thermo Fisher Scientific) and Sanger sequencing. Three clones with distinct out of frame indel formation were selected. Single clones with silent mutations provided phenotypic control cell lines. IDAA results were analysed using Peak Scanner Software V1.0 (Thermo Fisher Scientific). Human ST6GALNAC1 (hST6GALNAC1 [NM_018414.5]) knock-in (KI) was performed in *C1GALT1* KO cells and human S*T3GAL1* (hST3Gal1 [NM_173344.3]) KI was performed in wild-type and *GCNT1* KO cells by conventional mammalian gene expression vector transfection, using jetPRIME® transfection reagent (PolyPlus Transfection), according to manufacturer's instructions. Positively transfected cells were selected based on puromycin (2 µg/mL, EMD Millipore) resistance. In parallel, a mock system containing a 300 bp stuffer ORF was established.

**Autophagy assay**

Cells were cultured in 96 well plates and submitted to microenvironmental challenges of hypoxia and glucose deprivation. Non-challenged cells were used as control. The presence of autophagy was determined based on the detection of a proprietary fluorescent autophagosome marker using an Autophagy Assay Kit (MAK138, Sigma-Aldrich) according to the manufacturer's instructions. The fluorescence intensity was measured using a fluorescence microscope (Leica DMI6000 FFW).
