## Supplemental Figures for "Multilevel Plasticity and Altered Glycosylation Drive Aggressiveness in Hypoxic and Glucose-Deprived Bladder Cancer Cells"

### Supplementary Figures

Andreia Peixoto[1,2,3], Dylan Ferreira,[1,2,3#], Andreia Miranda[1,2,3#], Marta Relvas-Santos[1,2,3,4], Rui Freitas[1,2,3], Tim S. Veth[5,6], Andreia Brandão[1], Eduardo Ferreira[1], Paula Paulo[1], Marta Cardoso[1], Cristiana Gaiteiro[1,3], Sofia Cotton[1,3] Janine Soares[1,3,7], Luís Lima[1], Filipe Teixeira[8], Rita Ferreira[7], Carlos Palmeira[1,9,10], Albert J. R. Heck[5,6], Maria José Oliveira[2], André M. N. Silva[4], Lúcio Lara Santos[1,10,11], José Alexandre Ferreira[1,3*]

[1]Research Center of IPO-Porto (CI-IPOP) / RISE@CI-IPOP (Health Research Network), Portuguese Oncology Institute of Porto (IPO-Porto) / Porto Comprehensive Cancer Center (P.ccc) Raquel Seruca, Porto, Portugal; [2]i3S – Instituto de Investigação e Inovação em Saúde, Universidade do Porto, Porto, Portugal; [3]Institute of Biomedical Sciences Abel Salazar (ICBAS), University of Porto, Porto, Portugal; [4]LAQV-REQUIMTE, Department of Chemistry and Biochemistry, Faculty of Sciences, University of Porto, Porto, Portugal; [5]Biomolecular Mass Spectrometry and Proteomics, Bijvoet Center for Biomolecular Research and Utrecht Institute for Pharmaceutical Sciences, University of Utrecht, Utrecht, The Netherlands; [6]Netherlands Proteomics Center, Padualaan, Utrecht, The Netherlands; [7]QOPNA & LAQV-REQUIMTE, Department of Chemistry, University of Aveiro, Aveiro, Portugal; [8]Centre of Chemistry, University of Minho, Braga, Portugal; [9]Department of Immunology, Portuguese Oncology Institute of Porto, Porto, Portugal; [10]Health School of University Fernando Pessoa, Porto, Portugal; [11]Department of Surgical Oncology, Portuguese Oncology Institute of Porto, Porto, Portugal.

#equal contribution
*corresponding author

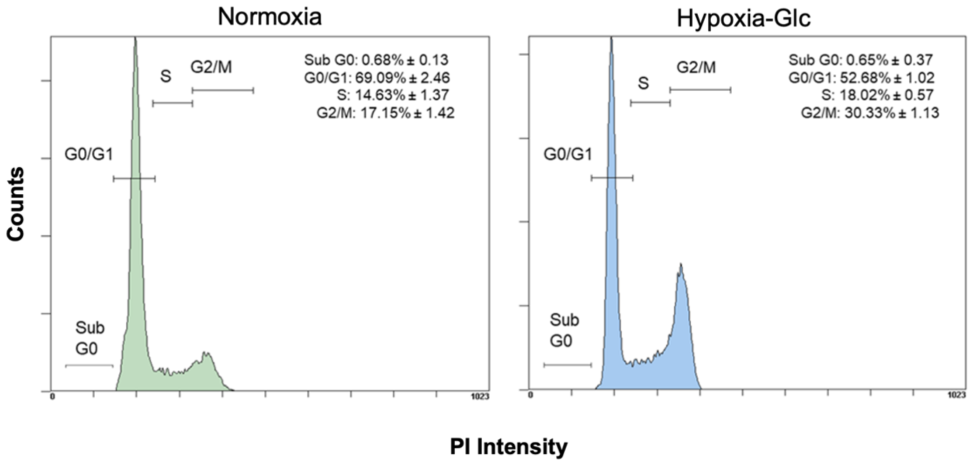


**Fig. 1. Hypoxia and low glucose induce an arrest of cell cycle in G2/M phases.** The histograms highlight the distribution of bladder cancer cells in normoxia and hypoxia and low glucose according to their cell cycle. This clearly shows an increase in the number of cells in G2/M under microenvironmental stress.

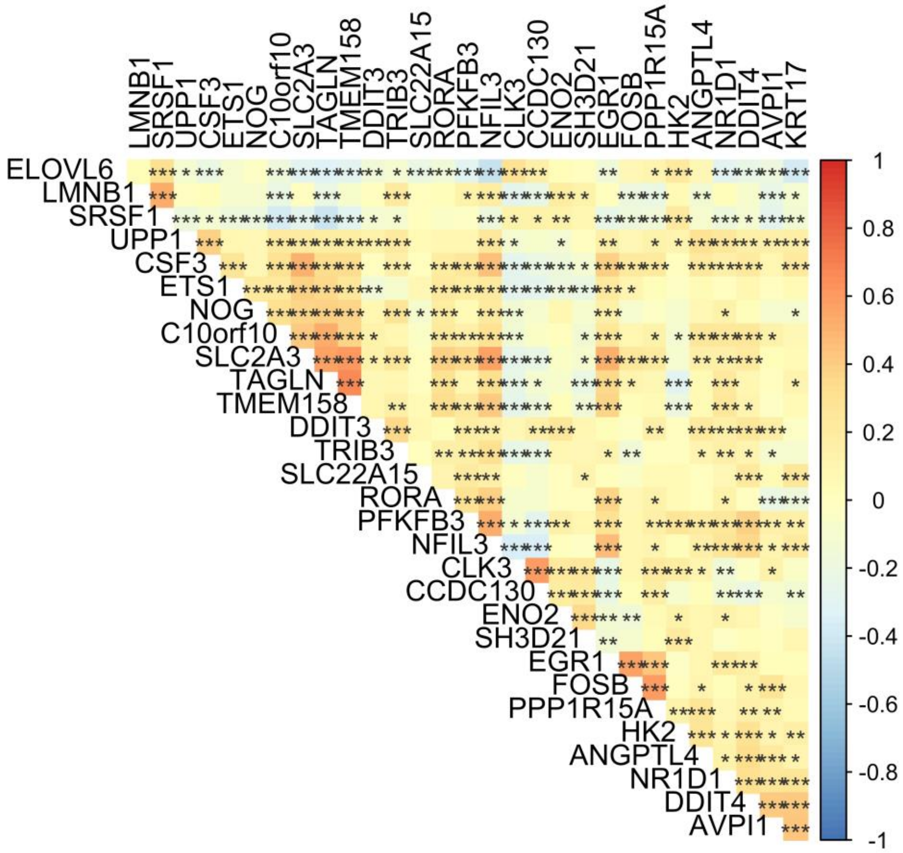


**Fig. 2. Differentially expressed genes in hypoxia and glucose-deprived cancer cells are also found in BLCA.** The correlogram for differentially expressed genes shows a high level of correlation between genes significantly linked to hypoxia and glucose deprivation in cancer cells also in BLCA tissue sections.

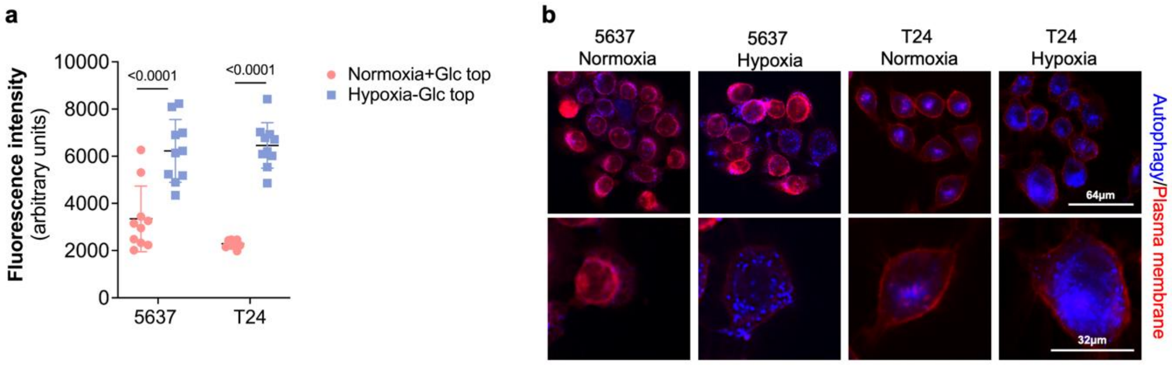


**Fig. 3. Hypoxic and glucose-deprived cells exhibit significantly higher levels of autophagy compared to non-stressed cells.** The cells demonstrate a higher percentage of autophagosomes in hypoxia and glucose deprivation, as compared to normoxia (**a and b**), as evident through fluorescence microscopy (**b**).

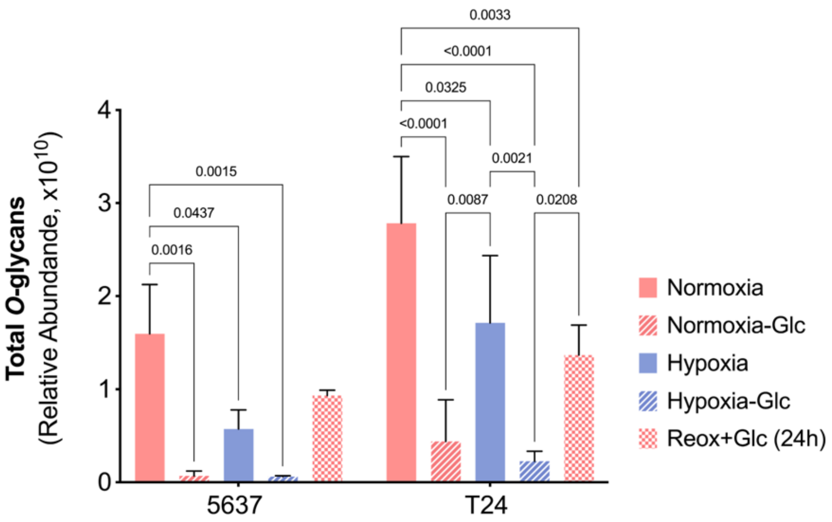


**Fig. 4. *O*-glycan biosynthesis is significantly inhibited by hypoxia and glucose deprivation, primarily driven by glucose suppression, and is partially restored after reoxygenation.** The suppression of glucose has a substantial impact on the efficiency of protein *O*-glycosylation, leading to significant consequences for the formation of glycosidic chains. Hypoxia has a comparatively lesser effect on this aspect, while the combined occurrence of both events results in *O*-glycans resembling those influenced by glucose. This strongly suggests that glucose suppression may be the primary influencing factor behind the inhibition of *O*-glycan biosynthesis. The biosynthesis capacity is significantly restored after 24 hours of reoxygenation.

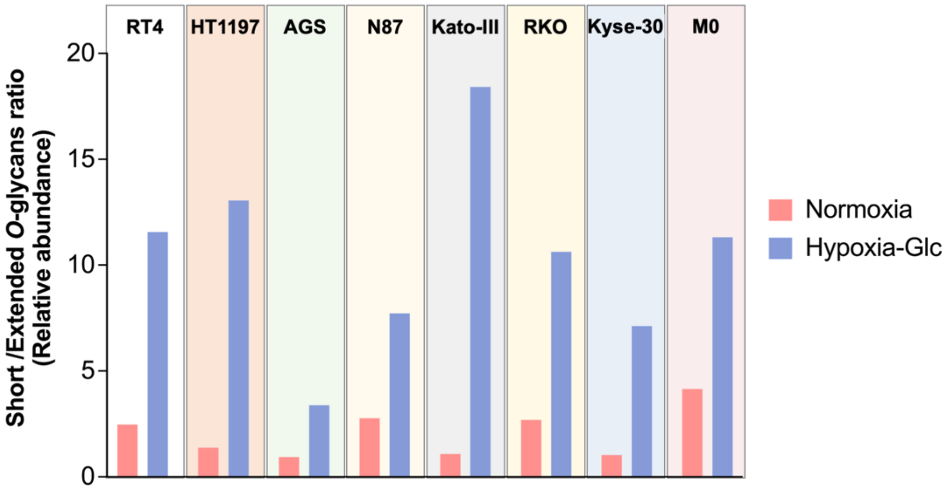


**Fig. 5. Hypoxia and glucose deprivation inhibit *O*-glycosylation extension across a wide range of different cell lines.** Cancer cell lines of different natures (RT4 and HT1197: bladder; AGS, N87 and Kato-III: gastric; RKO: colon; Kyse-30: oesophagus), as well as human monocyte-derived M0 macrophages, all exhibit a higher percentage of short-chain *O*-glycans in hypoxia compared to normoxia, when compared to glycosidic chains extended beyond core 2.

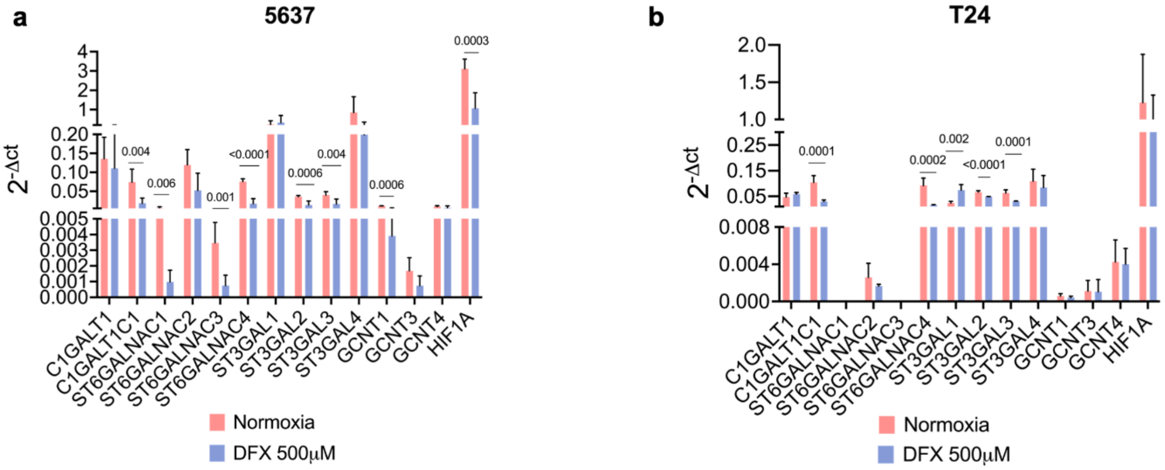


**Fig. 6. HIF-1α's influence on *O*-glycogenes expression varies depending on the cell line, consistently resulting in the downregulation of *C1GALT1C1*, *ST3GAL2*, *ST3GAL3*, and *ST6GALNAC4*.** Stabilization of HIF-1α with DFX induces significant changes in the expression of relevant *O*-glycogenes associated with elongation of glycosidic chains. This impact is more pronounced in 5637 cells (7 glycogenes significantly altered vs 5 in T24). Notably, all cell lines consistently exhibit reduced expressions of *C1GALT1C1*, a key factor for *O*-chains elongation beyond the Tn antigen. Also, *GCNT1*, responsible for core 2 biosynthesis, experiences downregulation in 5637 cells; in contrast, T24 cells display elevated *ST3GAL1* mRNA levels, which could also contribute to this premature termination. Notably, the consistent downregulation of *ST3GAL2*, *ST3GAL3*, and *ST6GALNAC4* triggered by HIF-1α supports a pivotal role for *ST3GAL1* in T sialylation.

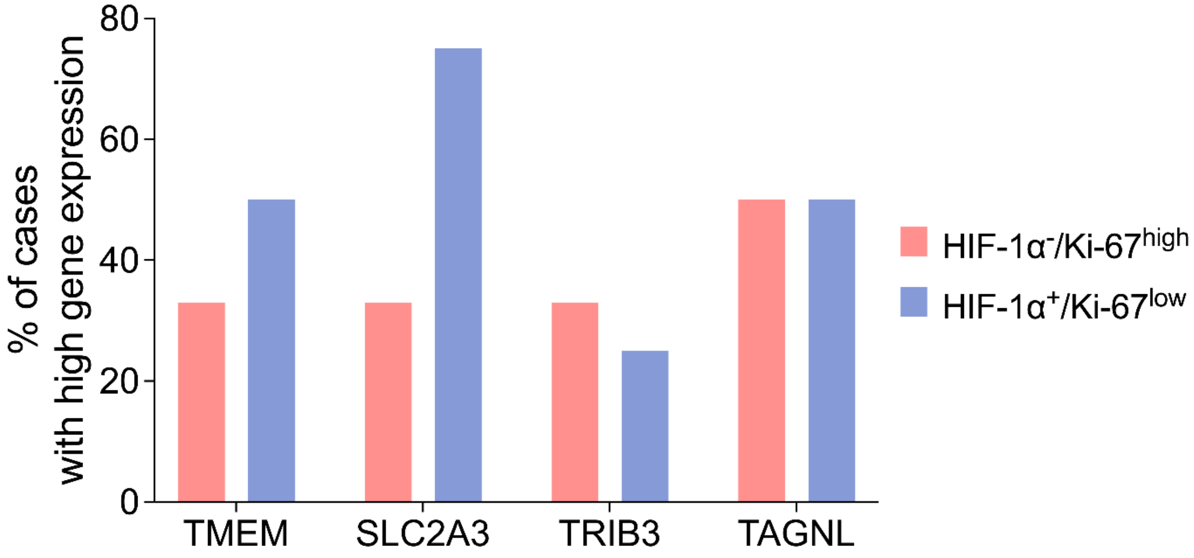


**Fig. 7. The frequency of tumours overexpressing hypoxic-linked poor prognosis transcripts *TMEM* and *SLC2A3* is higher for the HIF-1α$^{positive}$/Ki-67$^{low}$ compared to the HIF-1α$^{negative}$/Ki-67$^{high}$ group.** We screened a panel of transcripts associated with worse prognosis in BLCA under hypoxia and glucose deprivation conditions (TMEM, SLC2A3, TRIB3, TAGNL). We observed a higher frequency of tumours overexpressing TMEM and SLC2A3 in the HIF-1α$^{positive}$/Ki-67$^{low}$ group compared to the HIF-1α$^{negative}$/Ki-67$^{high}$ group. Notably, 75% of tumours in the HIF-1α$^{positive}$/Ki-67$^{low}$ group exhibited overexpression of two or more of these genes, in contrast to 33% in the HIF-1α$^{negative}$/Ki-67$^{high}$ group.

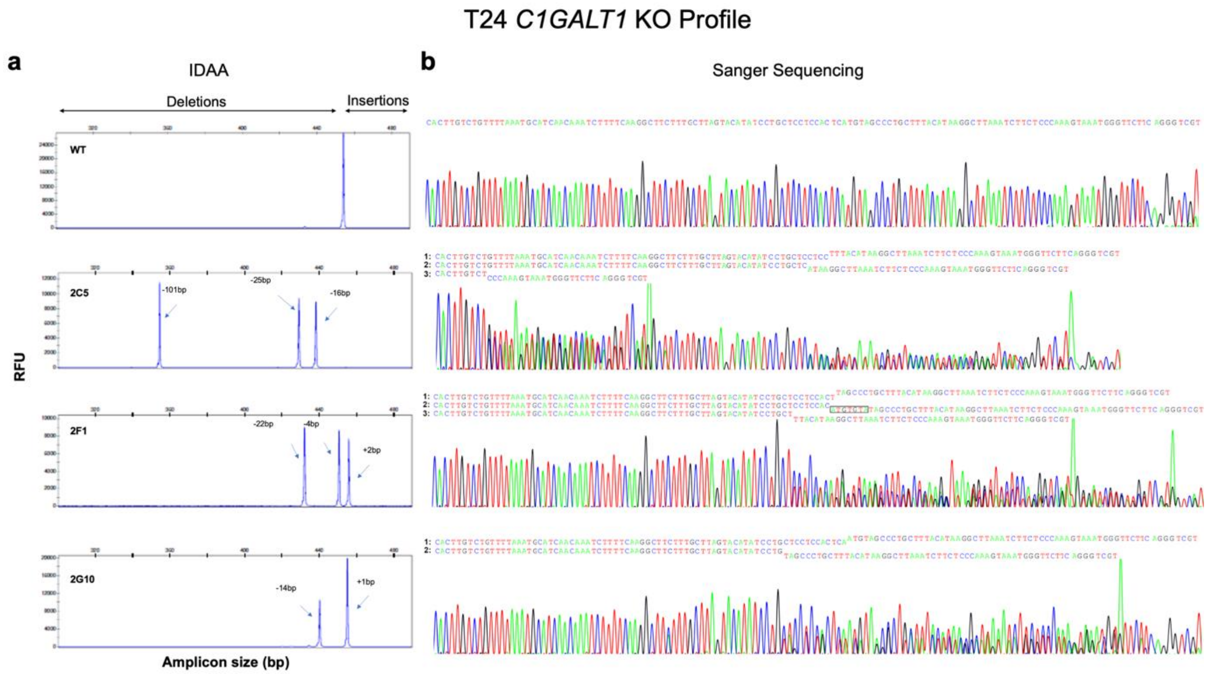


**Fig. 8. Genomic profiling of glycoengineered T24 *C1GALT1* KO models.** T24 cells were glycoengineered to knock-out human *C1GALT1* and three clones with diverse indels were selected for proof-of-concept experiments. Induced indel mutations were characterized by Indel Detection by Amplicon Analysis (IDAA) **(a)** and Sanger Sequencing **(b)**. Electropherograms of Sanger Sequencing with the reverse primer are shown, along with the corresponding sequencing readings (1, 2 and 3). Accordingly, the 2C5 clone is characterized by three different DNA sequences, with the following observed variants and predicted consequences: **1:** a c.610_625del [p.(Gln204GlufsTer7)], leading to a 16bp deletion and the replacement of Gln 204 by Glu, resulting in a frame-shift introducing a stop codon 7 a.a. ahead; **2:** a c.605_629del [p.(Val202GlufsTer6)], resulting in a 25bp deletion and the replacement of Val 202 by Glu, leading to a frame-shift introducing a stop codon 6 a.a. ahead; and **3:** a c.589_689del [p.(Arg197GlnfsTer5)], in which a 101bp deletion results in the replacement of Arg 197 by Gln, leading to a frame-shift introducing a stop codon 5 a.a. ahead. Similarly, clone 2F1 is also characterized by three different DNA sequences, with the following observed variants and predicted consequences: **1:** a c.618_621del [p.(Tyr206Ter)], where a 4bp deletion results in the replacement of Tyr 206 by a stop codon; **2:** a c.618_622delinsTACACAT [p.(Met207ThrfsTer10)], where a 5bp deletion and a 7bp insertion results in the replacement of Met 207 by Thr, inducing a frame-shift and introducing a stop codon 10 a.a. ahead; and **3:** a c.614_635del [p.(Gly205AspfsTer4)], where a 22bp deletion results in the replacement of Gly 205 by Asp, leading to a frame-shift introducing a stop codon 4 a.a. ahead. Finally, clone 2G10 is characterized by two different DNA sequences, with the following observed variants and predicted consequences: **1:** a c.620dup [p.(Met207IlefsTer19)], where the duplication of nucleotide 620 results in the replacement of Met 207 by Ile, leading to a frame-shift introducing a stop codon 19 a.a. ahead; and **2:** a c.620_633del [p.(Met207ArgfsTer14)], resulting in a 14bp deletion and the replacement of Met 207 by Arg, resulting in a frame-shift introducing a stop codon 14 a.a ahead.

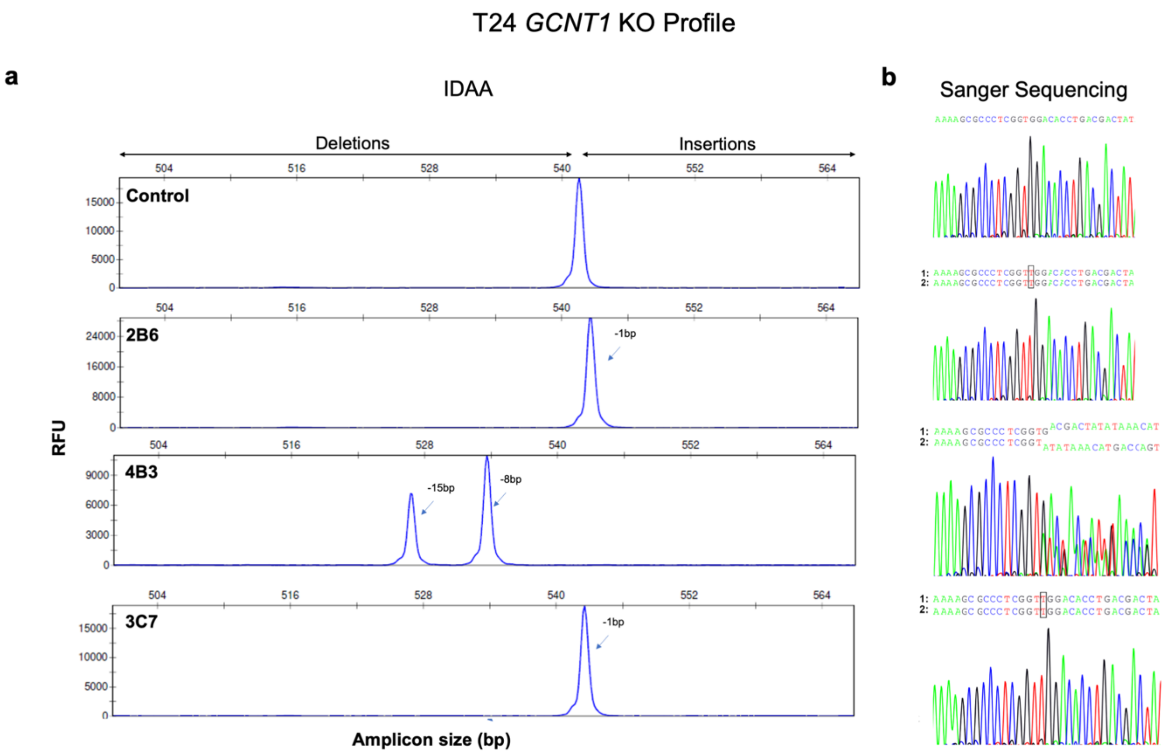


**Fig. 9. Genomic profiling of glycoengineered T24 *GCNT1* KO models.** T24 cells were glycoengineered to knock-out human *GCNT1* and three clones were selected for proof-of-concept experiments. Induced indel mutations were characterized by Indel Detection by Amplicon Analysis (IDAA) **(A)** and Sanger Sequencing **(B)**. Electropherograms of Sanger Sequencing with the forward primer are shown, along with the corresponding sequencing readings (1 and 2). Accordingly, both 3B6 and 3C7 where characterized by a single DNA sequence reading (assumed homozygous), with the c.262dup [p.(Trp88LeufsTer4)] being observed, where duplication of nucleotide 262 is predicted to lead to the replacement of Trp 88 by Leu, leading to a frame-shift introducing a stop codon 4 a.a ahead. In turn, 4B3 clone is characterized by two different DNA sequences, with the following observed variants and predicted consequences: **1:** a c.264_271del [p.(Trp88Ter)], where a 8bp deletion results in the replacement of Trp 88 by a stop codon; and **2:** a c.263_277del [p.(Trp88_Asp92del)], where a 15bp in-frame deletion between Trp 88 and Asp 92 results in the suppression of 5 a.a and the generation of a truncated protein.

a

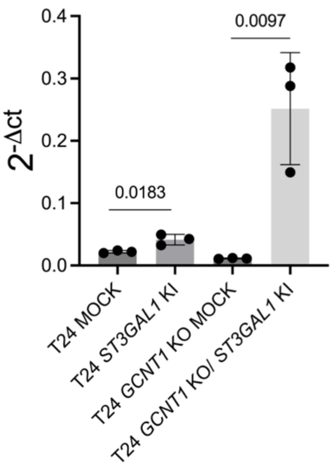


**Fig. 10. *ST3GAL1* knock-in on T24 and T24 *GCNT1* KO cells results in a significant increase in the expression of this glycosyltransferase.**

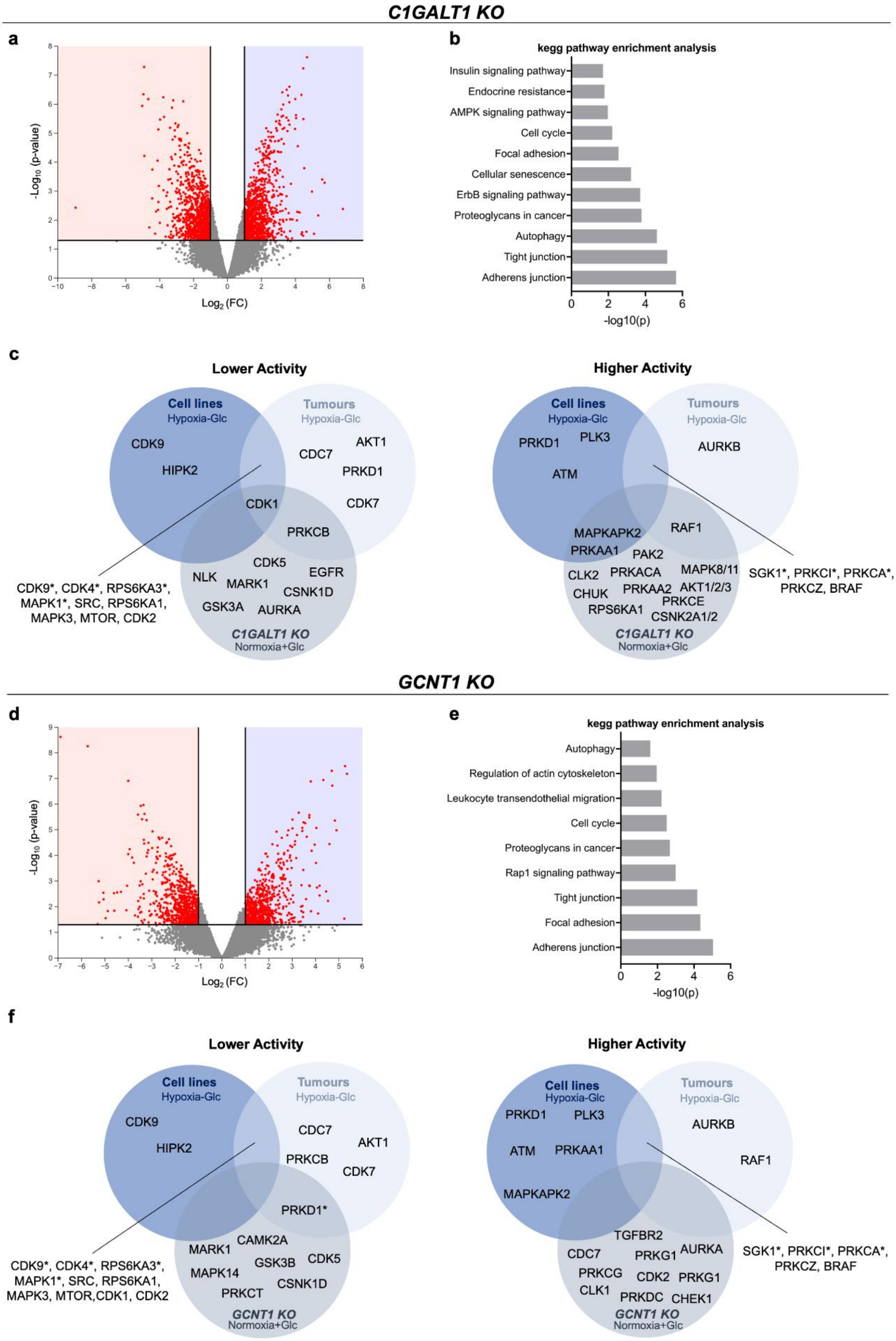


**Fig. 11. Induction of Simple O-Glycophenotypes Drives Profound Molecular Changes in Signaling Networks Supporting Cancer Aggressiveness. a.** Suppression of O-glycan elongation beyond the Tn antigen (T24 C1GALT1 KO) leads to significant and substantial alterations in cellular signaling networks, highlighting

the impact of glycosylation changes on cancer aggressiveness. **b.** KEGG pathway enrichment analysis of phosphoproteomics data reveals significant modifications in cell signalling pathways associated with cell-cell and cell-matrix adhesion, autophagy, and cellular senescence—patterns also observed in stressed cells. **c.** T24 C1GALT1 KO cells exhibit distinct kinase activity profiles compared to stress-exposed T24 wild-type cells and hypoxic tumours. Predicted activation or inhibition is indicated by z-scores ≥2 or ≤ -2, with significance at $p < 0.05$. Kinases with $p < 0.07$ are denoted by *. **d.** Suppression of O-glycan elongation beyond core 1 (T24 GCNT1 KO) has a comparatively milder impact on protein phosphorylation compared to C1GALT1 KOs. **e.** KEGG pathway enrichment analysis of phosphoproteomics data also demonstrates significant alterations in cell signaling pathways, supporting the shared influence of glycosylation on cellular functions. **f.** T24 GCNT1 KO cells exhibit distinct kinase activity profiles compared to stress-exposed wild-type cells and tumors, emphasizing the crucial role of microenvironmental pressure in cellular signaling rewiring. For **c** and **f**, predicted activation or inhibition is indicated by z-scores ≥2 or ≤ -2, with significance at $p < 0.05$. Kinases with $p < 0.07$ are denoted by *.
